# Benchmarking single-cell multi-modal data integrations

**DOI:** 10.1101/2025.04.01.646578

**Authors:** Shaliu Fu, Shuguang Wang, Duanmiao Si, Gaoyang Li, Yawei Gao, Qi Liu

## Abstract

Single-cell multi-modal omics, remarked as “Method of the Year 2019” by Nature Methods, provides valuable insights into complex tissues. Recent advances have enabled the generation both unpaired (separate profiling) and paired (simultaneous measurement) multi-modal datasets, driving rapid development of single-cell multi-modal integration tools. Nevertheless, there is a pressing need for a comprehensive benchmark to assess algorithms under varying integrated dataset types, integrated modalities, dataset sizes, and data quality. Here, we present a systematic benchmark for 40 single-cell multi-modal integration algorithms involving modalities of DNA, RNA, protein and spatial omics for paired, unpaired and mosaic datasets (a mixture of paired and unpaired datasets). We evaluated usability, accuracy, and robustness to assist researchers in selecting suitable integration methods tailored to their datasets and applications. Our benchmark provides valuable guidance in the ever-evolving field of single-cell multi-omics.

The Stage 1 protocol for this Registered Report was accepted in principle on 3rd Jul 2024. The protocol, as accepted by the journal, can be found at https://springernature.figshare.com/articles/journal_contribution/Benchmarking_single-cell_multi-modal_data_integrations/26789572.

## Main

Single-cell multi-modal omics has rapidly developed in recent years^1^, enabling profiling of DNA, RNA, and protein at the single-cell level. Current single-cell multi-modal datasets can be broadly categorized into two groups based on the relationship between different modalities^2^. The first category involves unpaired single-cell multi-modal sequencing, where cells are segregated into distinct batches for individual omics profiling. The second category is paired single-cell multi-modal sequencing, which allows simultaneous measurement of various modalities within the same cell. Moreover, single-cell multi-modal sequencing can be further classified based on the targeted modalities, each with significantly varying dimensions^3^. For instance, it can detect thousands of genes for mRNA, hundreds of thousands of chromatin accessible regions for DNA, but only tens to hundreds of antibody-derived tags (ADT) for proteins. In addition, spatial multiomics can further capture subcellular resolution along with gene expression, chromatin accessibility^4^ or protein abundance^5^.

To gain a deeper understanding of the interplay between multiple omics in single cells, bioinformaticians have developed several types of integration methods tailored to specific single-cell multi-modal integration tasks. These multi-modal integration methods can be classified into three major types: paired integration methods for paired multi-modal data from the same cells, unpaired diagonal integration methods for unpaired multi-modal data, and unpaired mosaic integration methods, which integrate both paired and unpaired multi-modal data with a shared modality type. Additionally, these integration methods can be further classified by the types of modalities used for integration.

An essential challenge in the context of single-cell multi-modal integration is how to effectively leverage the heterogeneity inherent in different modalities. Firstly, the dimensions of various modalities can vary significantly, spanning from just a handful of proteins to hundreds of thousands of chromatin accessible sites. Secondly, the characteristics of different modalities can also be highly diverse. For instance, a single gene in scRNA data can be detected with several number of reads in each cell, while the chromatin accessibility site in scATAC data is often binary, representing either an open or closed state. Moreover, scATAC data exhibits a much greater degree of data sparsity, with sequencing reads distributed across hundreds of thousands of chromatin-accessible sites, in contrast to scRNA data, where sequencing reads are only concentrated in thousands of genes.

To address these challenges in single-cell multi-modal integration, researchers have developed over 40 computational tools for this task. A prior study analyzed five tools for single-cell RNA and ATAC integration and four Multiome-guided mosaic integration tools^6^. However, there remains a need for a comprehensive benchmark to evaluate algorithms across diverse dataset types, modalities, sizes, and qualities. In this study, we conducted a systematic benchmark for paired and unpaired diagonal and unpaired mosaic single-cell multi-modal integration methods concerning the modalities of chromatin accessibility (DNA), gene expression (RNA), ADT (protein) and spatial omics. For each multi-modal integration type, we assessed the usability, accuracy and robustness for each integration algorithm. We also developed a user-friendly platform for accessing comprehensive benchmark results, and a benchmark pipeline that will empower researchers to assess and choose optimal integration methods for their specific dataset processing requirements, as well as the evaluation of new integration algorithms.

## Results

### Single-cell multi-modal integration benchmark (SCMMIB)

To systematically evaluate single-cell multi-modal integration tools, we collected 40 algorithms from published or preprint papers. These methods include bindSC^7^, CiteFuse^8^, cobolt^9^, DCCA^10^, DeepMAPS^11^, GCN-SC^12^, GLUE^13^, Liger iNMF^14^, Liger UiNMF^15^, Liger online iNMF^16^, MaxFuse^17^, MEFISTO^18^, MIDAS^19^, MOFA+^20^, Multigrate^21^, MultiMAP^22^, MultiVI^23^, Pamona^24^, SAILERX^25^, scAI^26^, SCALEX^27^, scDEC^28^, sciPENN^29^, scMCs^30^, scMDC^31^, scMM^32^, scMoMaT^33^, scMVAE^34^, scMVP^35^, scVAEIT^36^, Seurat v3 CCA^37^, Seurat v4 RPCA^37^, Seurat v4 WNN^38^, Seurat v5 bridge^39^, SIMBA^40^, SpatialGlue^41^, StabMap^42^, totalVI^43^, unionCom^44^, uniPort^45^. On the basis of the single-cell modalities (DNA, RNA, protein, spatial omics) and integration types (paired, unpaired diagonal, unpaired mosaic) used, we categorized these methods by six specific integration types, including 17 paired scRNA and scATAC integration methods, 13 paired scRNA and ADT integration methods, 3 spatial multiomics integration methods, 14 unpaired scRNA and scATAC diagonal integration methods, 8 scRNA and scATAC mosaic integration methods and 8 scRNA and ADT mosaic integration methods (Figure 1). In summary, 40 multi-modal integration algorithms were benchmarked as 65 integration methods in total, as some algorithms support multiple integration tasks (Supplementary Table 1, Supplementary Notes 1-2).

**Fig. 1.**
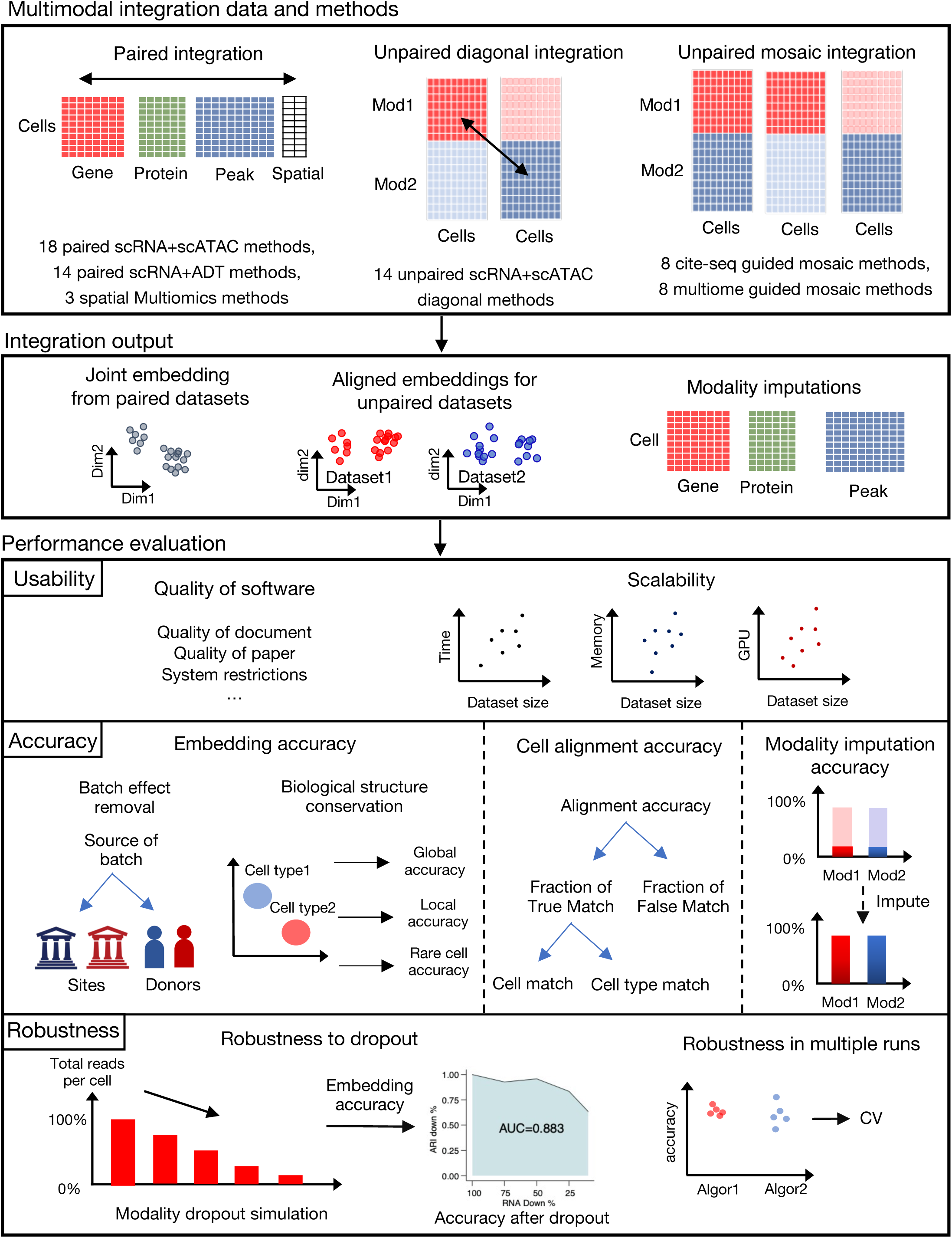
Illustration of single-cell multi-modal integration benchmark (SCMMIB) study. We collected 65 single-cell multimodal integration methods from 40 algorithms to evaluate their performance across paired integration tasks, unpaired diagonal integration tasks, and unpaired mosaic integration tasks. The integration outputs—joint embeddings from paired multimodal datasets, aligned embeddings from unpaired multimodal datasets, and modality imputations—were generated for evaluating usability, accuracy, and robustness. Each benchmark method was assessed on these three criteria to provide a comprehensive performance evaluation.

To perform a comprehensive evaluation of these multi-modal integration methods, we collected 14 publicly available single-cell multiomics datasets to generate 101 benchmark datasets for these multi-modal integration tasks (Supplementary Table 2), which covered various cell types (such as immune cells, kidney, skin, and brain), dataset sizes ranging from 1,000 to 700,000 cells and batch counts ranging from 2 to 143 (Extended Data Figs. 1-3, Supplementary Figs. 1-3). Specifically, all these datasets were applied for accuracy evaluations, and datasets with curated and unbiased cell annotations were used for robustness evaluations (details in Methods). For the BMMC Multiome and CITE-seq datasets, we further refined the level 1 cell types on the basis of the original level 2 cell type annotations to assess the accuracy metrics at different levels of granularity (Supplementary Table 3).

For all integration tasks, we established a unified benchmark workflow to evaluate the usability, accuracy, and robustness of integration methods, as illustrated in Fig. 1:

(1) For the usability assessment, we considered the user-friendliness of the method, encompassing the quality of the associated paper, code, and documentation for each method. Additionally, we assessed whether the algorithm could run within reasonable and pre-defined computational hardware limits across variable sizes of multi-omics datasets.
(2) For the accuracy assessment, we applied different evaluation metrics for paired and unpaired multi-modal integration. For paired integration algorithms, the focus was put on jointly embedding different modalities with known cell-cell matches into a common latent space, and this was evaluated for the accuracy of the single joint embedding (embedding accuracy). For unpaired diagonal integration algorithm, it aims to match similar cells from different modalities, and their accuracy was evaluated for alignment accuracy across embeddings of integrated modalities (cell alignment accuracy) as well as embedding accuracy of separate modalities after integration. The unpaired mosaic integration method aims to integrate paired and unpaired multi-modal datasets into the same space, which is evaluated for both embedding accuracy for the common embedding of paired and unpaired datasets, as well as cell alignment accuracy in the unpaired datasets. We used paired multi-modal datasets as pseudo-unpaired multi-modal datasets for the evaluation of unpaired diagonal or mosaic integration methods, which omitted all or part of the cell-matching information from different modalities. We then took the cell match information as the gold standard for pseudo-unpaired multi-modal datasets. The embedding accuracy was evaluated with metrics related to biological structure conservation and batch effect removal (Methods). The cell alignment accuracy was evaluated for the false cell match fraction and the true cell match fraction at the cell and cell type levels on pseudo unpaired multi-modal datasets. For paired and mosaic integration methods with imputation function, we also assessed whether the posterior imputation can effectively mitigate data sparsity in simulated high dropout datasets or impute missing modalities in pseudo mosaic datasets.
(3) For the robustness assessment, we evaluated the accuracy of algorithms when confronted with multi-modal datasets featuring varying degrees of data sparsity in either a single modality or both modalities. To begin, we identified optimal accuracy metrics for assessing robustness. Robustness was quantified by evaluating the extent of accuracy decline via a multi-gradient AUC metric (Methods) and by assessing the accuracy metrics in the highest dropout datasets. For unpaired mosaic integration, we also evaluated the robustness to paired multi-modal dataset sizes (3,000 to 20,000 cells) with the multi-gradient AUC metric. Additionally, we assessed the stability of the accuracy of the integration methods by performing five independent runs.

### Performance of paired scRNA and scATAC integration methods

We first performed benchmark analysis of methods for multimodal datasets with known cell matches in scRNA and scATAC modalities, also known as paired scRNA and scATAC integration methods. For paired scRNA and scATAC integration, we evaluated usability metrics (Fig. 2a). Most methods (13/16) support GPU acceleration, with MultiVI and uniPort performing best in usability. Among these benchmark methods, MultiVI, uniPort, and scMVP were available for BMMC Multiome simulation datasets of 500,000 cells. scAI was not available for datasets of 50,000 cells because of excessively long computation time, while DCCA and scMVAE were only available for 5,000 and 10,000 cells because they exceeded the GPU memory limit (Supplementary Table 4). uniPort took longer to run on smaller datasets because it set more training epochs for smaller datasets. Among these integration methods, MOFA+ and DeepMAPS support the biological interpretability of the latent embedding. MOFA+ uses feature weights in each factor of feature embeddings to explain the relative importance of latent factors in cell latent space, while DeepMAPS uses attention mechanisms in a heterogeneous graph transformer model to describe relationships between genes and cells.

**Fig. 2.**
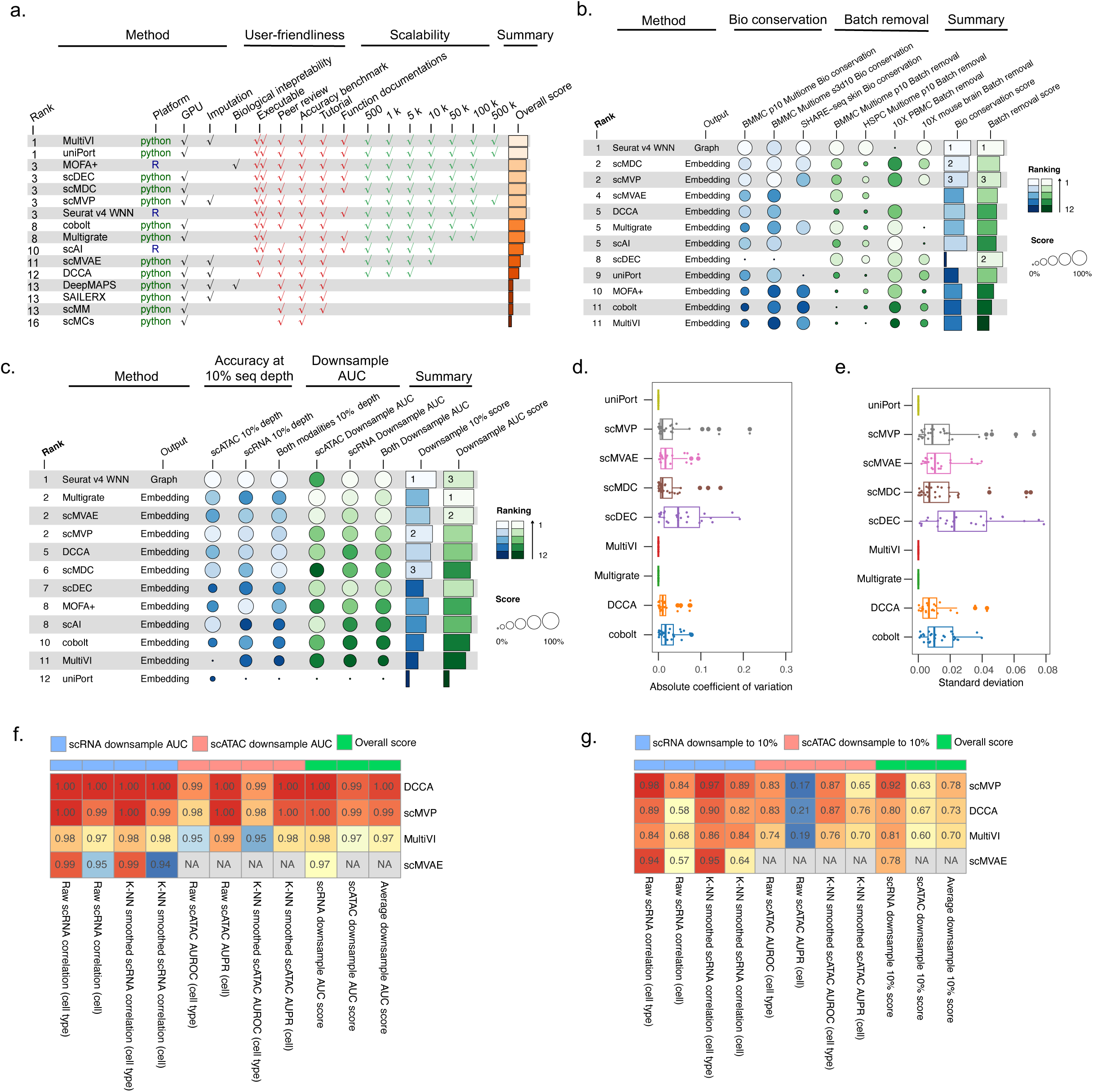
Benchmark results for paired scRNA and scATAC integration methods. **a**, Overview of usability results, including user-friendliness metrics and scalability metrics. For the executable metric in user-friendliness metrics, methods can be directly executed from source code scored 2 (“ √√ ” in figure), methods that required debugging to run received scored 1 (“√” in figure), and nonexecutable methods scored 0. For scalability metrics, methods that completed the task within 500GB of memory and 24 hours were scored as 1 (“√” in figure), while others scored as 0. The overall score was the sum of all usability metrics. **b**, Overview of accuracy results, including biological conservation and batch removal metrics. The rank of each benchmark method was determined by averaging the ranks of the summary scores for the biological conservation and batch removal metrics. **c,** Overview of robustness results, including downsample AUC metrics (Methods) and accuracy at the 10% downsampled sequencing depth. Method ranks were computed as average ranks of summary scores for downsample AUC and accuracy at the 10% downsampled sequencing depth. **d**-**e**, Robustness of accuracy metrics across 5 repeated runs, quantified by **(d)** absolute coefficient of variation (CV) and **(e)** standard deviation (SD). Box plots show the median (center line), the 25th and 75th percentiles (bounds of the box), and whiskers extend to 1.5 × IQR (interquartile range). **f**-**g**, Accuracy of paired scRNA and scATAC imputation in recovering true signal from downsampled datasets. Imputed profiles from downsampled data were evaluated using either the raw profile or k-NN smoothed profile at the cell or cell type level, measured by Pearson correlation, AUROC, or AUPRC (Methods). Performance was assessed using (**f)** the downsample AUC score and (**g)** accuracy at the 10% downsampled sequencing depth ..

For paired scRNA and scATAC integration methods that could be executed, we evaluated the accuracy via both biological conservation metrics and batch removal metrics (Fig. 2b, Extended Data Fig. 4b, Supplementary Figs. 4-5). Batch information was not provided as input for any benchmark methods to ensure a fair comparison. For the biological conservation metrics, Seurat v4, scMDC, and scMVP were ranked as the top three methods (Fig. 2b). Seurat v4 WNN achieved high performance in the BMMC p10 dataset with mixed batches and two single batch datasets. scMDC also showed high accuracy in two single batch datasets, but was inferior to Seurat v4 WNN in the BMMC p10 mixed batches dataset (Extended Data Fig. 4b). For the batch removal metrics, Seurat v4, scDEC, and scMVP achieved the best performance among the paired scRNA and scATAC integration algorithms. However, the ranks of the algorithms were inconsistent across different benchmark datasets. For example, Seurat v4 WNN showed the highest performance in BMMC p10, HSPC p10, and 10X mouse brain datasets, but was inferior to the other algorithms on the 10X PBMC dataset. The inconsistency in rankings across different datasets might be due to the limited number of batch removal metrics available for datasets without well-curated cell annotations, for example, only graph iLISI for the graph output method, such as Seurat v4 WNN (Supplementary Fig. 5).

We further evaluated the influence of the batch parameter on the accuracy of the methods via the BMMC Multiome p10 dataset. Among the paired scRNA and scATAC integration methods, Multigrate and MultiVI allow batch information as an optional input parameter. We found that providing batch information improved 4 out of 5 batch removal metric scores in MultiVI, and 5 out of 8 biological conservation metric scores in both MultiVI and Multigrate (Extended Data Fig. 4c). These improvements in accuracy metrics indicate the necessity of batch information input for these two algorithms.

Next, we evaluated the robustness of paired scRNA and scATAC integration algorithms to data sparsity through downsampling simulations (Methods). To identify optimal accuracy metrics for the robustness evaluation, we compared biological conservation metrics and batch removal metrics in all methods at different downsampling levels (Supplementary Fig. 6). The biological conservation metrics were heavily affected by downsampling levels, showing high correlations across different downsampling levels in the scRNA and multimodal downsampling simulations (Supplementary Fig. 7a-b). Next, we used the biological conservation metric scores at the lowest downsampling levels and the downsample AUC of these metrics to evaluate the robustness of the paired scRNA and scATAC integration algorithms (Methods). Overall, Seurat v4 WNN, Multigrate, scMVAE and scMVP ranked highest according to both metrics. For the biological conservation metrics at the 10% downsampling level, Seurat v4 WNN, scMVP and scMDC achieved the best performance (Fig. 2c, Extended Data Fig. 4d, Supplementary Fig. 7c), similar with their high biological conservation scores in the accuracy evaluations. For the downsample AUC metrics, Multigrate, scMVAE, and Seurat v4 WNN presented a high overall score (Fig. 2d, Extended Data Fig. 4d), and scMVP presented an AUC over 0.95 for all metrics (Supplementary Fig. 7d). These findings suggested the high robustness of top integration methods to data sparsity problems in paired scRNA and scATAC datasets.

To evaluate the impact of the random sampling step on method accuracy, we further assessed the robustness of the output with standard deviation (SD) and the absolute coefficient of variation (CV) of the accuracy metric scores for each method (Fig. 2d, e). All non-deep learning methods generated exactly same output in repeated runs. Among the deep learning methods, uniPort, MultiVI and Multigrate generated the exact same output in five runs by setting fixed random seeds in the source code. scDEC showed highest coefficient of variations and a median standard deviation of 0.02 for all metrics, and scMVP and scMDC also suffered from high variations in some accuracy metrics.

To assess the imputation accuracy of the four algorithms with imputation function, we first evaluated the nonzero ratio of the ground truth data at the cell and cell type levels (Extended Data Fig. 4e, Supplementary Table 5). For both the scRNA and scATAC modalities, k-NN smoothing mitigated the data sparsity in each cell. Then, we selected the optimal metric for the scATAC modality evaluation. The nonzero ratios in the scATAC data were much lower than 0.5 at the cell level, indicating far fewer positive labels than negative labels in the accuracy assessment and indicating that AUPR (Area Under Precision Recall curve) was the optimal metric. Similarly, as nonzero ratios were greater than 0.5 at the cell type level, we selected the AUROC (Area Under the Receiver Operating Characteristic) as the optimal metric for scATAC imputation accuracy at the cell-type level. We next evaluated the imputation accuracy at all downsampling levels via the same downsample AUC metrics and the accuracy at the lowest sequencing depth in a robustness evaluation. For the four benchmark methods, the imputations of DCCA and scMVP were merely influenced by the downsampling levels (Fig. 2f). scMVP showed the best imputation accuracy for the scRNA modality, while DCCA performed best for scATAC modality imputation at both the cell and cell type levels (Fig. 2g).

### Performance of paired scRNA and ADT integration methods

In addition to paired scRNA and scATAC integration methods, some methods integrate scRNA with low-dimensional modalities like ADT in paired cells.. To assess the performance of the paired scRNA and ADT integration methods, we first evaluated the usability of the benchmark methods (Fig. 3a, Extended Data Fig. 5a). MaxFuse, MOFA+, SCALEX, scMDC, and totalVI met all the user-friendliness and scalability criteria. In scalability tasks, Seurat v4 WNN and bindSC exceeded the 500 GB memory limit on simulation datasets with 500,000 cells. We also encountered gradient explosion issues when running Multigrate on datasets with 100,000 and 500,000 cells, as well as an index error in CiteFuse with datasets of 50,000 cells or more (Supplementary Table 6). In addition to MOFA+, totalVI also enables biological interpretation, utilizing an archetype analysis neural network to identify biologically meaningful archetypes in the nonlinear latent space of cells. scMM and DeepMAPS were excluded from accuracy and robustness evaluations because of limitations in code usability (Methods).

**Fig. 3.**
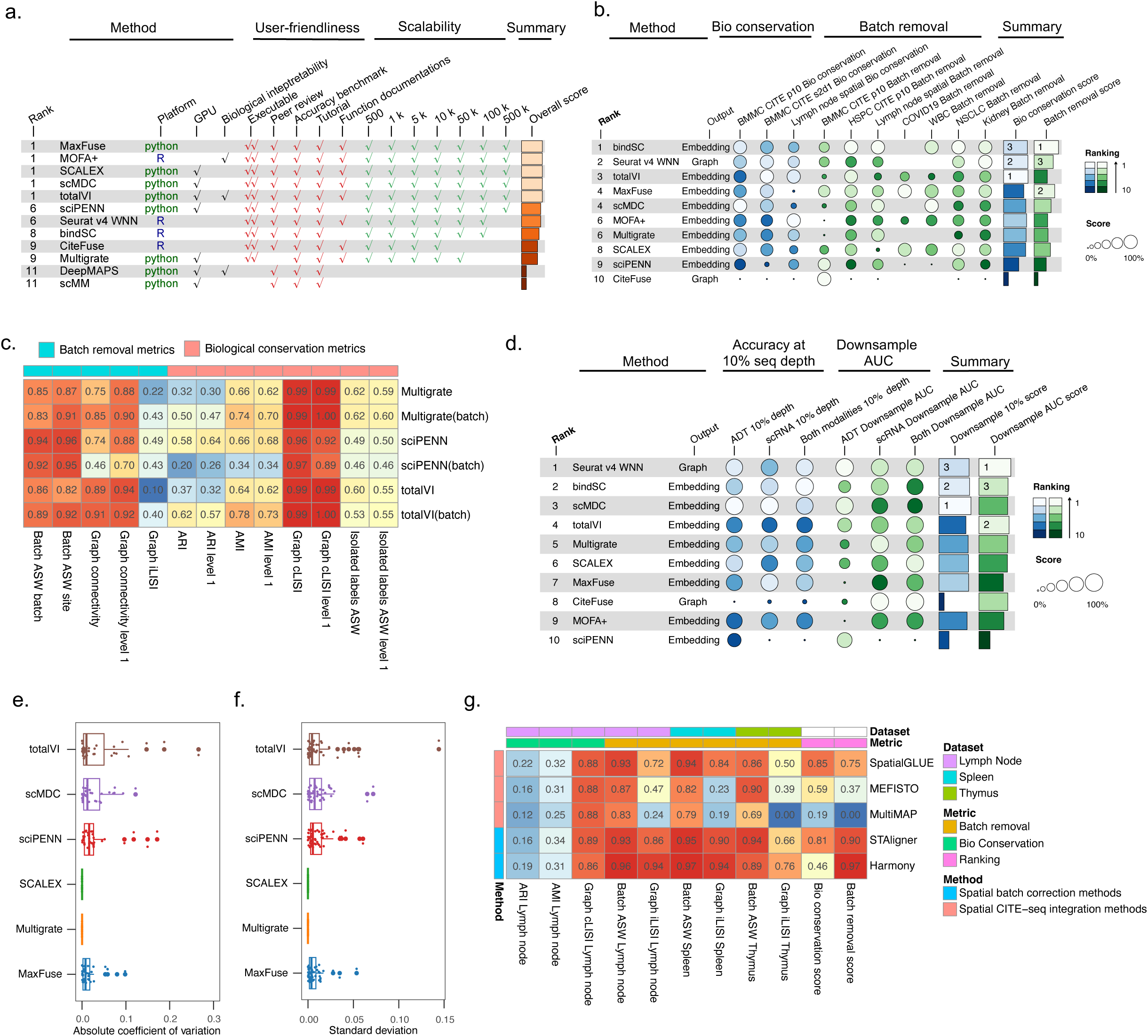
Benchmark results for paired scRNA and ADT integration methods. **a**, Overview of usability results, including user-friendliness metrics and scalability metrics. **b**, Overview of accuracy results, including metrics of biological conservation and batch removal. The rank of the benchmark method was computed as the average of the ranks for the biological conservation summary score and batch removal summary score. **c,** Evaluation for algorithms with optional batch parameter. The metric results calculated with batch information are labeled with the “(batch)” suffix in figure. **d,** Overview of robustness results, including downsample AUC metrics and accuracy at the 10% downsampled sequencing depth. The rank of benchmark method was computed as the average of the ranks for downsample AUC summary score and accuracy at the 10% downsampled sequencing depth summary score. **e-f**. Stability of algorithm embedding output in 5 repeated runs, evaluated with (**e**) absolute coefficient of variation and (**f**) standard deviation of metric values for all accuracy metrics. Box plots show the median (centre line), the 25th and 75th percentiles (bounds of the box), and whiskers extend to 1.5 × IQR (interquartile range). **g,** Accuracy metrics of the spatial scRNA and ADT integration methods compared with those of two spatial scRNA data batch correction methods.

Next, we evaluated the accuracy of the executable paired scRNA and ADT integration methods. Overall, totalVI, Seurat v4 WNN, and bindSC showed the best performance in terms of the biological conservation metrics (Fig. 3b, Supplementary Figs. 8-9). totalVI ranked in the top two for two single-batch datasets but only 8th out of 10 methods in the mixed-batch dataset, highlighting the significant impact of batches on the totalVI’s biological conservation accuracy without batch information input (Extended Data Fig. 5b). bindSC, scMDC, and Seurat v4 WNN performed best in terms of batch removal metrics. bindSC demonstrated consistently high performance across all executable datasets, ranking 4th in the BMMC p10 dataset and in the top two for the other five benchmark datasets (Extended Data Fig. 5b).

We investigated the influence of the batch parameter on the accuracy of paired scRNA and ADT integration algorithms using the BMMC CITE-seq p10 dataset. For both totalVI and Multigrate, providing batch information significantly improved the biological conservation and batch removal metrics of both algorithms. For example, the batch removal metric graph iLISI increased from 0.1 to 0.4 for totalVI and from 0.22 to 0.43 for Multigrate, second only to sciPENN among all paired scRNA and ADT integration methods (Fig. 3c). The biological conservation metric ARI also increased from 0.37 to 0.62 for totalVI and from 0.32 to 0.50 for Multigrate. However, the inclusion of the batch parameter did not improve the accuracy of sciPENN.

To assess the robustness of the paired scRNA and ADT integration methods, we conducted the same accuracy metric evaluations as those for the paired scRNA and scATAC methods and compared biological conservation and batch removal metrics across all the methods at various downsampling levels. The levels of modality sparsity were positively correlated with biological conservation metrics but had less of an impact on metric scores than the paired scRNA and scATAC methods did (Supplementary Figs. 10 and 11a-b), as the ADT modality experienced fewer data sparsity issues in downsampling simulations (Extended Data Figs. 1-2). We then evaluated the robustness of the algorithms using both biological conservation metrics at the 10% sequencing depth and downsample AUC metrics (Fig. 3d, Extended Data Fig. 5c, Supplementary Fig. 11c-d). scMDC, bindSC, and Seurat v4 WNN performed best at the 10% downsampling level. Seurat v4 WNN, totalVI, and bindSC demonstrated the highest downsample AUC across all downsampling levels.

We also assessed the robustness of the paired scRNA and ADT integration methods across repeated runs. All non-deep learning methods consistently generated stable outputs. The metrics for the two deep generative models, SCALEX and Multigrate, were also stable across five repeats, because of the use of a fixed random seed in the source code (Fig. 3e-f). The performance of the other four algorithms fluctuated across different runs, with standard deviations exceeding 0.05 in certain metrics.

Recent advances in single-cell multiomics technologies and bioinformatics methods have incorporated the paired scRNA and ADT sequencing with spatial omics^41, 46, 47^. Therefore, we further evaluated the performance of three available methods for spatial multiomics integration, along with two commonly used batch correction methods for spatial-omics (Fig. 3g). Among the three spatial multiomics integration methods, SpatialGlue, which is specifically designed for spatial multiomics, outperformed the other two methods across all accuracy metrics. SpatialGlue also outperformed the two spatial batch correction methods, Harmony^48^ and STAligner^49^, in terms of biological conservation metrics, but was inferior to these two methods in terms of batch removal metrics. STAligner demonstrated high accuracy in both biological conservation and batch removal metrics when using only spatial location and scRNA modality as inputs in spatial CITE-seq data, indicating the importance of batch correction for current spatial multiomics integration methods applied to spatial CITE-seq datasets.

### Performance of unpaired diagonal integration methods

Several single-cell multimodal integration algorithms have been developed for scRNA and scATAC datasets that lack known cell match information between the two modalities. These methods are referred to as unpaired scRNA and scATAC diagonal integration methods (Fig. 1). We first assessed the usability of these algorithms (Fig. 4a, Extended Data Fig. 6a). Unlike paired multimodal integration methods that use direct modality input, the majority (9 out of 14) of the unpaired scRNA and scATAC integration methods require an independently computed gene activity matrix (GAM) from the scATAC modality as input (Supplementary Table 7). The GAM is crucial for aligning the two unpaired modalities through genes shared between the GAM and the scRNA modality.

**Fig. 4.**
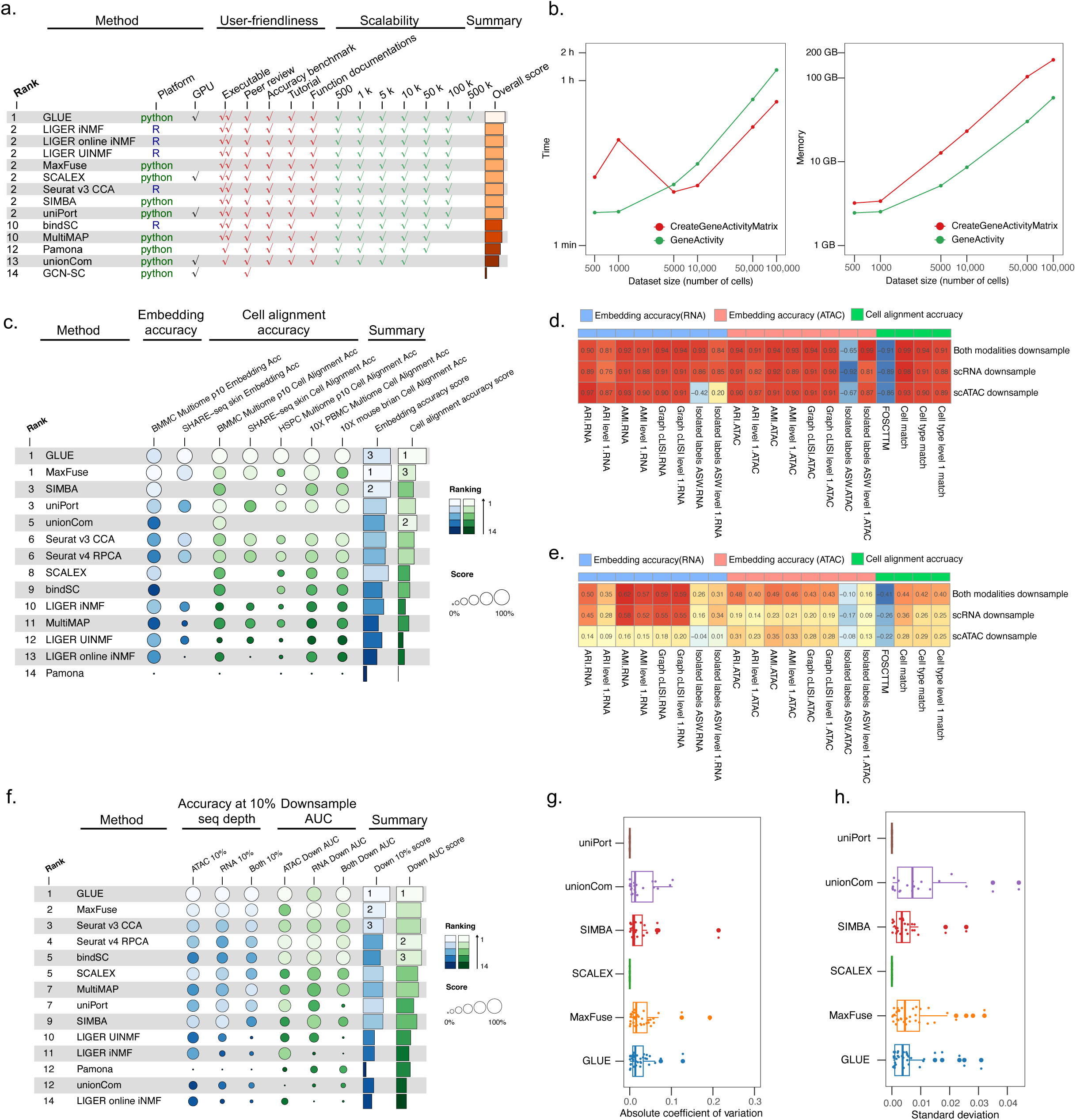
Benchmark results for unpaired scRNA and scATAC diagonal integration methods. **a**, Overview of usability results, including user-friendliness metrics and scalability metrics. **b,** The scalability of two gene activity matrix (GAM) functions (GeneActivity and CreateGeneActivityMatrix) for preparing unpaired scRNA and scATAC diagonal integration input data. **c**, Overview of accuracy results, including metrics of embedding accuracy and cell alignment accuracy. The rank of the benchmark method was computed as the average of the ranks for the embedding accuracy summary score and the cell alignment accuracy summary score. **d-e**, Correlations between metric values and downsampling levels. The color and number in heatmaps indicate the Pearson’s r correlation between (**d)** metric values or (**e**) average metric values with the downsampling levels. **f,** Overview of robustness results, including downsample AUC metrics and accuracy at the 10% downsampled sequencing depth. The rank of the benchmark method was computed as the average of the ranks for the downsample AUC summary score and accuracy at the 10% downsampled sequencing depth summary score. **g-h**. Stability of algorithm embedding output in 5 repeated runs, evaluated with (**g**) absolute coefficient of variation and (**h**) standard deviation of metric values for all accuracy metrics. Box plots show the median (center line), the 25th and 75th percentiles (bounds of the box), and whiskers extend to 1.5 × IQR (interquartile range).

To evaluate the scalability of GAM preparation, we tested two common functions from the Signac^50^ and Seurat packages^37^. The GeneActivity function in the Signac package, which estimates gene activity from the entire gene body via scATAC fragment files, is the most recent method for computing GAMs. In contrast, the CreateGeneActivityMatrix function, which is applicable only in earlier versions of the Seurat package^37^, computes a GAM on the basis of gene promoter regions via scATAC accessible matrix data. Unfortunately, neither GAM preparation function was compatible with the BMMC Multiome simulation dataset containing 500,000 cells, thereby limiting the scalability of methods relying on GAM input for this large dataset (Fig. 4b). Specifically, the GeneActivity function processed the GAM as a sparse matrix, which exceeded R’s object length limit for the 500,000-cell dataset. Moreover, the CreateGeneActivityMatrix function surpassed the scalability resource limit for this same dataset (Supplementary Table 8). Consequently, GLUE emerged as the only algorithm capable of handling the 500,000-cell dataset, demonstrating the highest usability for unpaired scRNA and scATAC diagonal integration (Fig. 4a). In contrast, GCN-SC exhibited limited usability metrics and was excluded from accuracy and robustness evaluations.

To assess the accuracy of the executable unpaired scRNA and scATAC diagonal integration methods, we evaluated both cell alignment and embedding accuracy post integration for the unpaired modalities (Fig. 4c). In terms of embedding accuracy, MaxFuse, SIMBA, and GLUE performed best in the BMMC Multiome p10 dataset, with GLUE and MaxFuse achieving higher embedding accuracy scores than the other methods did in the SHARE-seq skin dataset (Extended Data Fig. 6b). In terms of cell alignment accuracy, GLUE outperformed all the other metrics across all the datasets, indicating a significant advantage in this area (Supplementary Figs. 12-13). UnionCom and MaxFuse ranked just behind GLUE in cell alignment accuracy; however, UnionCom was only applicable to only one benchmark dataset due to its limited usability (Extended Data Fig. 6b).

To identify optimal accuracy metrics for robustness analysis, we evaluated the impact of downsampling levels on both the embedding accuracy and the cell alignment accuracy metrics. The scores for these metrics were significantly affected by the level of data sparsity (Supplementary Fig. 14). The downsampling levels of the scRNA, scATAC, or both modalities strongly correlated with the accuracy of the cell alignment metrics. Notably, compared to the scATAC modality, the downsampling levels of the scRNA modality and both modalities had greater impacts on the embedding accuracy metrics (Fig. 4d-e). We subsequently assessed the robustness of the unpaired scRNA and scATAC diagonal integration methods by embedding accuracy and cell alignment accuracy metrics (Fig. 4f, Extended Data Fig. 6c, Supplementary Fig. 15). GLUE performed the best at a 10% downsampling depth, while MaxFuse and Seurat v3 CCA also demonstrated high performance for this metric. Across all downsampling levels, GLUE, Seurat v4 RPCA, and bindSC achieved the highest downsample AUC scores for robustness. We also examined the robustness of these methods through five repeated runs (Fig. 4g-h). Among the non-deep-learning methods, SIMBA and MaxFuse display fluctuations in metric scores and output results across these repetitions. In contrast, the deep learning-based methods SCALEX and uniPort showed stability in different runs, owing to the fixed random seed setting. Overall, GLUE showed the highest performance in usability, accuracy, and robustness to low sequencing depth.

### Unpaired scRNA and scATAC mosaic integration benchmark

To evaluate the performance of single-cell multimodal integration methods for partially paired scRNA and scATAC datasets, also referred to as unpaired scRNA and scATAC mosaic integration methods, we first assessed their usability on the basis of user-friendliness and scalability metrics. For this purpose, we used BMMC Multiome simulation datasets, which included both paired scRNA and scATAC data as well as pseudo-unpaired datasets of varying sizes (Methods). Overall, MultiVI, Seurat v5 Bridge, and StabMap satisfied all the evaluation criteria and ranked the highest (Fig. 5a, Supplementary Table 9). In contrast, MIDAS exceeded both the time and memory limits for the combined dataset of 20,000 paired and 20,000 unpaired cells (denoted as c20k_c20k in Extended Data Fig. 7a). Similarly, scMoMaT and scVAEIT exceeded the GPU memory limits for large-scale datasets. Notably, scVAEIT was excluded from accuracy and robustness evaluations, as it could only process datasets with only 1,000 paired and 1,000 unpaired cells—significantly smaller than the datasets used in our accuracy and robustness benchmarks (Extended Data Fig. 3).

**Fig. 5.**
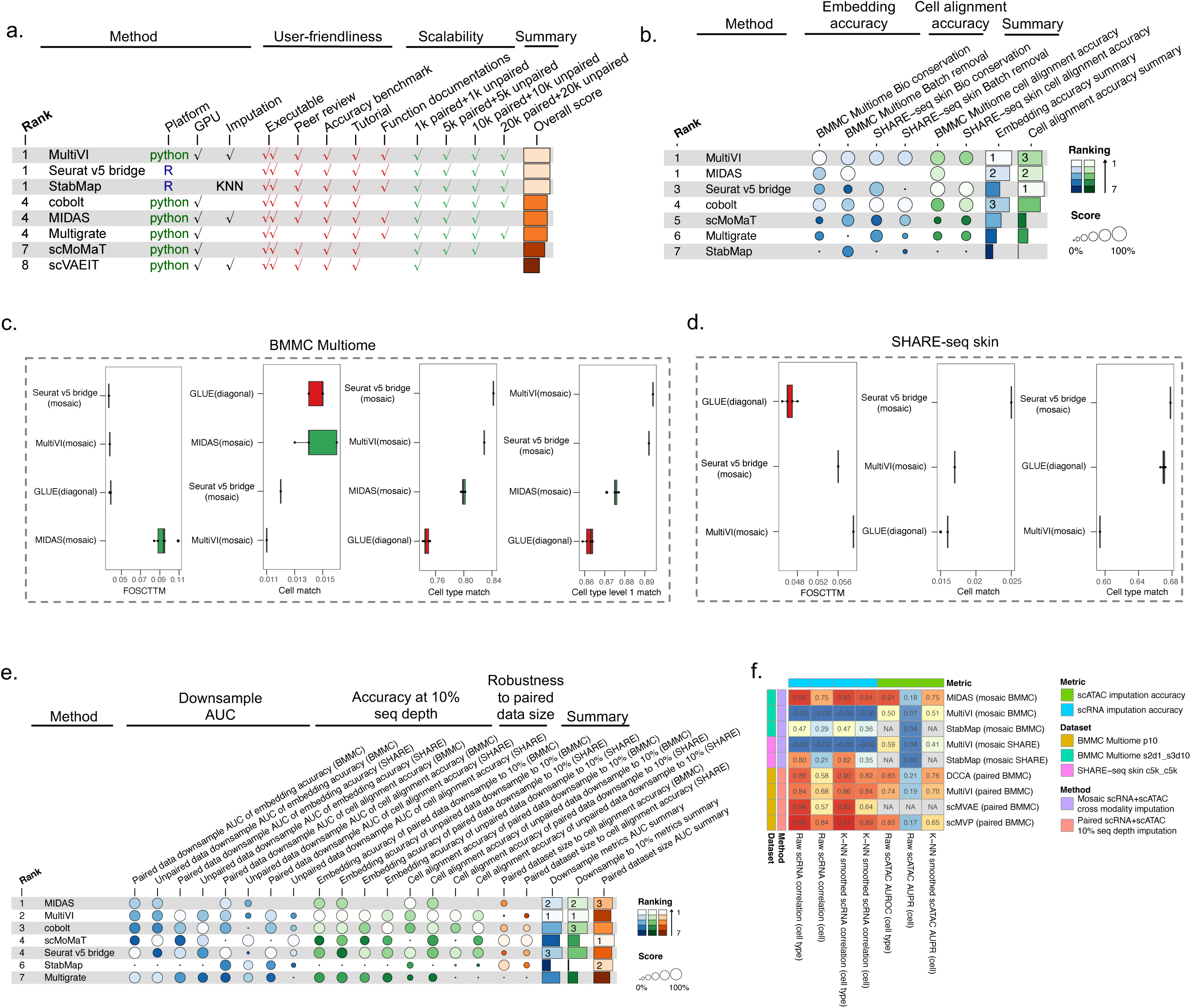
Benchmark results for unpaired scRNA and scATAC mosaic integration methods. **a**, Overview of usability results, including user-friendliness metrics and scalability metrics. **b**, Overview of accuracy results, including metrics of the embedding accuracy (including biological conservation metrics and batch removal metrics) and the cell alignment accuracy. The rank of the benchmark method was computed as the average of the ranks for the embedding accuracy summary score and the cell alignment accuracy summary score. **c**-**d**, The boxplot of cell alignment accuracy metrics for the top three unpaired scRNA and scATAC mosaic integration methods and the unpaired diagonal integration method GLUE across 5 repeated runs, which were applied to the (**c**) BMMC Multiome dataset and (**d**) SHARE-seq skin dataset. Box plots show the median (centre line), the 25th and 75th percentiles (bounds of the box), and whiskers extend to 1.5 × IQR (interquartile range). **e,** Overview of robustness results in BMMC and SHARE-seq skin mosaic simulation datasets, including downsample AUC metrics and accuracy at the 10% downsampled sequencing depth for both paired and unpaired datasets downsample, as well as AUC metrics for paired dataset size. The rank of benchmark method was computed as the average of ranks for the paired multimodal dataset size AUC summary score, downsample AUC summary score and accuracy at the 10% downsampled sequencing depth summary score. **f,** The accuracy of cross-modality imputation (predicting scATAC from scRNA, and vice versa) for the unpaired scRNA and scATAC mosaic integration methods, compared with paired scRNA and scATAC imputation accuracy at the 10% sequencing depth downsampled datasets. The imputation accuracy was assessed both at the cell level and at the cell type level for each metric.

To assess the accuracy of the unpaired scRNA and scATAC mosaic integration methods, we compared the embedding accuracies of various benchmark methods on the BMMC Multiome and SHARE-seq simulation datasets (Fig. 5b, Extended Data Fig. 7b, Supplementary Figs. 16-17). The embedding accuracy for each algorithm was calculated via cells composed of three sub-datasets: paired scRNA and scATAC data, unpaired scRNA data, and unpaired scATAC data. The objective was to assess the degree of integration among these datasets by treating the three sub-datasets as different batches. Among the methods, MultiVI, MIDAS, and cobolt exhibited the best embedding accuracy across all cells. Seurat v5 Bridge, MIDAS, and MultiVI also performed well in aligning unpaired cells. However, MIDAS could not be applied to the SHARE-seq dataset due to excessive computational time and memory requirements when processing the high-dimensional (344,592 peaks) scATAC data (Extended Data Fig. 3).

Compared with unpaired diagonal integration, unpaired mosaic integration methods appear to integrate unpaired cells more effectively by leveraging paired multiome cells. To test this hypothesis, we next compared the cell alignment accuracy of the top three unpaired scRNA and scATAC mosaic integration methods with that of GLUE, a leading unpaired diagonal integration method (Fig. 5c). Seurat v5 Bridge and MultiVI outperformed GLUE in FOSCTTM (median of 3.7% and 3.8% versus 3.9%) and cell type matching metrics at both level 1 (median of 89.2% and 89.4% versus 86.3%) and level 2 (84.2% and 82.9% versus 74.5%) annotations. However, GLUE achieved the highest cell match score (median of 1.5%) in the BMMC Multiome dataset. Seurat v5

Bridge also surpassed GLUE in terms of cell match (median of 2.5% versus 1.6%) and cell type match (median of 67.8% versus 67.0%), although GLUE performed better in FOSCTTM than Seurat v5 Bridge did (median of 4.6% versus 5.6% and 5.9%) (Fig. 5d). These findings suggested an advantage in using paired scRNA and scATAC datasets with similar cell types to align unpaired cells via leading unpaired scRNA and scATAC mosaic integration methods.

To evaluate the robustness of the unpaired scRNA and scATAC mosaic integration methods, we applied downsampling simulations to both paired and unpaired datasets, as well as to the number of cells in paired multiome data, which has been reported as a crucial factor for the accuracy of the unpaired scRNA and scATAC mosaic integration^6^. We used three categories of metrics to assess robustness: downsample AUC, accuracy at the 10% downsampling level for both embedding accuracy across all cells, and cell alignment accuracy in unpaired cells. Additionally, we evaluated the robustness of the methods concerning the sizes of the paired datasets (Methods). Across all embedding accuracy and cell alignment accuracy metrics, biological conservation metrics and cell alignment accuracy metrics showed consistent changes in robustness, which were used to evaluate the overall robustness of the methods (Supplementary Fig. 18). MultiVI ranked highest on the downsample AUC metrics and accuracy at the 10% downsampling level, demonstrating superior performance in handling dataset sparsity (Fig. 5e, Extended Data Fig. 7c, Supplementary Fig. 19). scMoMaT, StabMap, and MIDAS also performed well in terms of robustness, particularly with respect to paired dataset sizes. MIDAS ranked within the top three for all robustness metrics in the BMMC dataset, indicating a high robustness to both dataset sizes and sequencing quality.

Some unpaired scRNA and scATAC mosaic integration methods support cross-modality imputation for unpaired modalities. For instance, StabMap employs a k-nearest neighbors (k-NN) approach to impute the missing modality via paired cells with both modalities. Some deep generative methods impute the missing modality from latent embeddings, similar to approaches used in paired scRNA and scATAC integration methods for downsampled datasets. We applied the same evaluation metrics for the scRNA and scATAC modalities as those used in paired scRNA and scATAC imputation, but excluded the metric that uses k-NN smoothed scATAC as the ground truth, as the nonzero ratio for MultiVI imputation was 99.99% (Supplementary Table 5). Since StabMap considers only the top scATAC peaks for imputation and the nonzero ratio at the cell type level is 1, we evaluated StabMap using the raw data as the ground truth at the cell level. We subsequently evaluated the cross-modality imputation accuracy of the unpaired scRNA and scATAC mosaic integration methods and compared it to the imputation accuracy of the paired scRNA and scATAC integration methods (Fig. 5f). MIDAS demonstrated significantly higher imputation accuracy for both the scRNA and scATAC modalities than did MultiVI and k-NN imputation in StabMap. Notably, the cross-modality imputation accuracy of MIDAS was comparable to that of four paired scRNA and scATAC imputation methods, which were provided with 10% of the original sequencing data in the target modality.

### Unpaired scRNA and ADT mosaic integration benchmark

To assess the performance of the unpaired scRNA and ADT mosaic integration methods, we first evaluated the usability of the benchmark methods (Fig. 6a, Extended Data Fig. 8a, Supplementary Table 10). Given the substantially lower dimensionality of the ADT modality (∼100 features) than the scATAC modality (>100,000 features), all eight benchmark methods were applicable to the scalability datasets. However, the computational resource consumption varied significantly among the methods. For instance, MIDAS required over 10 hours and 80 GB of memory to process the largest scalability dataset, whereas StabMap completed the same task in approximately two minutes using only 5 GB of memory. Additionally, scVAEIT consumes over 22 GB of GPU memory for all scalability datasets, which is near the GPU memory limit in our benchmark.

**Fig. 6.**
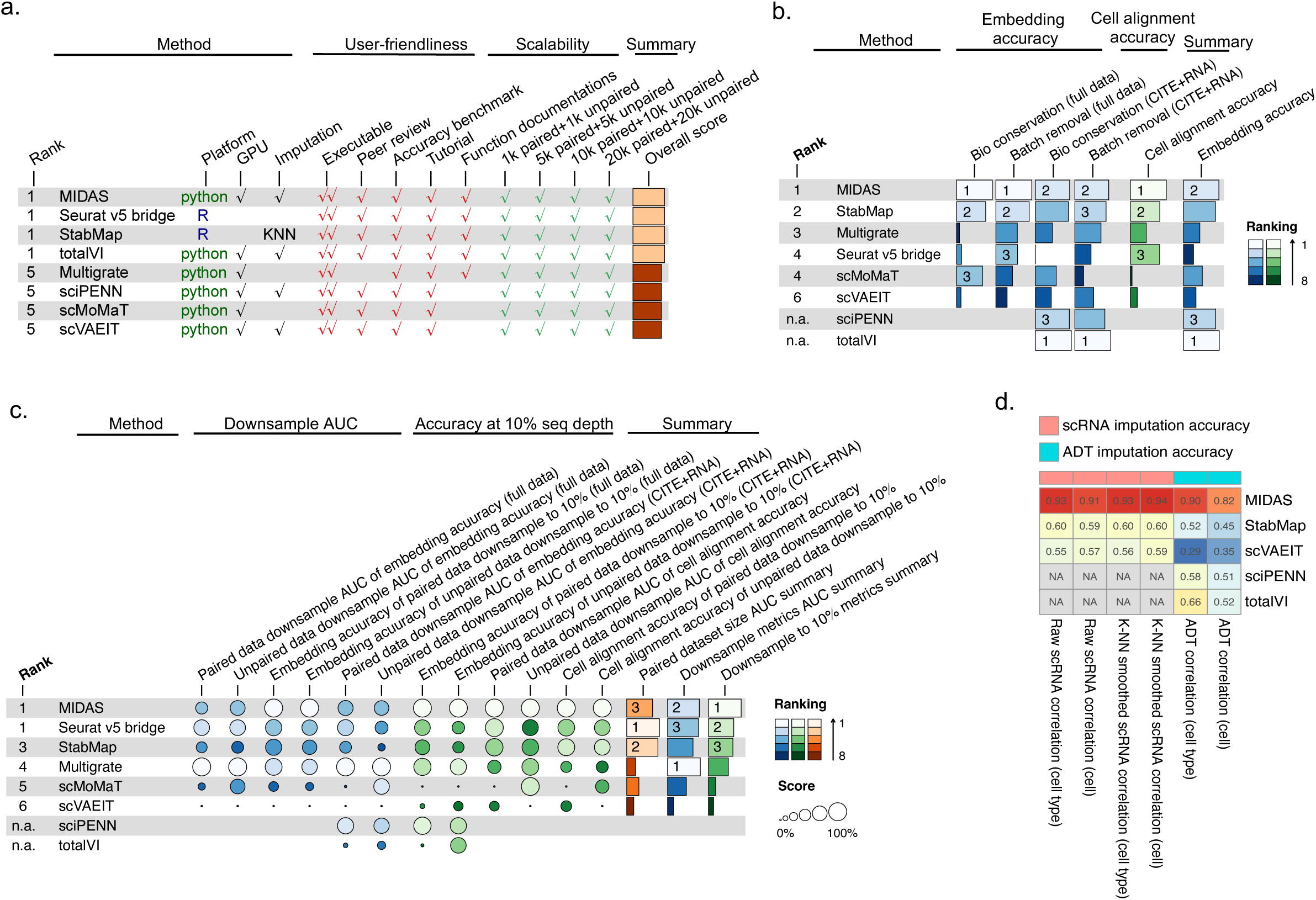
Benchmark results for the unpaired scRNA and ADT mosaic integration methods. **A**, Overview of usability results, including user-friendliness metrics and scalability metrics. **b**, Overview of accuracy results, including metrics of embedding accuracy (including biological conservation metrics and batch removal metrics) and cell alignment accuracy. As sciPENN and totalVI support only CITE-seq and scRNA integration, we evaluated the embedding accuracy of benchmark methods on the CITE-seq and scRNA dataset (referred to as “CITE+RNA” in the figure) as well as the full scRNA and ADT mosaic dataset (referred to as “full data” in the figure). The rank of each benchmark method was computed as the average of the ranks for the embedding accuracy summary score and the cell alignment accuracy summary score. **c,** Overview of robustness results for the full mosaic integration dataset as well as the CITE-seq and scRNA dataset, including downsample AUC metrics, accuracy at the 10% downsampled sequencing depth, and AUC metrics evaluating the effect of paired dataset size on unpaired integration performance. The rank of the benchmark method was the average of the for the paired dataset size AUC summary score, downsample AUC summary score and accuracy at the 10% downsampled sequencing depth summary score. **d,** The accuracy of cross modality imputation (predicting ADT from scRNA, and vice versa) for unpaired scRNA and ADT mosaic integration methods. The imputation accuracy was assessed both at the cell level and at the cell type level for each metric.

Since some unpaired scRNA and ADT mosaic integration methods only support integration of the paired scRNA and ADT datasets with unpaired scRNA data, we conducted evaluations on both the full mosaic scRNA and ADT simulation dataset, which was used for algorithms supporting full integration, and on the CITE-seq and scRNA datasets, which were applicable to all benchmark methods (Fig. 6b, Extended Data Fig. 8b, Supplementary Figs. 19-20). In the complete BMMC CITE-seq mosaic integration dataset, MIDAS and StabMap ranked highest in terms of embedding accuracy according to the biological conservation and batch effect removal metrics. Additionally, MIDAS, StabMap, and Seurat v5 bridge achieved the best performance in terms of cell alignment accuracy. For the CITE-seq and scRNA dataset, totalVI and MIDAS excelled in both biological conservation and batch effect removal metrics, while StabMap and sciPENN also demonstrated strong performance in embedding accuracy (Extended Data Fig. 8b).

To evaluate the robustness of the unpaired scRNA and ADT mosaic integration methods, we assessed their accuracy at varying levels of data sparsity and different sizes of the paired scRNA and ADT datasets. All unpaired scRNA and ADT mosaic integration methods were affected by data sparsity levels in both the paired and unpaired datasets, as well as by the size of the paired datasets (Supplementary Fig. 21). Overall, Seurat v5 bridge, StabMap, and MIDAS exhibited greater robustness to changes in paired dataset sizes than the other methods did (Fig. 6c, Extended Data Fig. 8c, Supplementary Fig. 22). MIDAS and Seurat v5 bridge performed best on the downsample AUC metric, with MIDAS showing a notable lead in accuracy metrics at the 10% downsampling level. We also evaluated the robustness of the methods across repeated runs (Extended Data Fig. 8d-e). Multigrate and scMoMaT consistently generated the same outputs in repeated runs by setting a random seed. However, MIDAS and scVAEIT presented greater variance than sciPENN and totalVI did, with standard deviations exceeding 0.05 for a few metrics. These findings highlighted the exceptional performance of MIDAS in terms of both accuracy and robustness for unpaired scRNA and ADT mosaic integration.

Additionally, we assessed the cross-modality imputation accuracy of five methods that support imputation (Fig. 6d). Similar to the exceptional performance observed in the unpaired scRNA and scATAC mosaic integration methods, MIDAS substantially outperformed in imputation accuracy across all scRNA and ADT evaluation metrics among unpaired scRNA and ADT mosaic integration methods, including totalVI. Notably, totalVI was reported as the best method for protein abundance prediction among 14 benchmark methods in a recent study^51^, which did not include MIDAS.

## Discussion

In this study, we developed a unified benchmark analysis workflow for six tasks of single-cell multimodal integration and conducted a comprehensive evaluation of 65 integration methods across 40 algorithms (Fig. 1). For each multimodal integration task, we identified top-performing methods across different metrics. Notably, two algorithms—Seurat v4 WNN and GLUE—consistently outperformed others in their respective tasks, establishing themselves as state-of-the-art (SOTA) methods for paired scRNA and scATAC integration, and unpaired scRNA and scATAC diagonal integration, respectively (Extended Data Fig. 9). Furthermore, we identified MIDAS as the SOTA method for cross-modality imputation in both unpaired mosaic integration tasks, significantly surpassing other imputation methods and approaching the accuracy of the paired scRNA and scATAC imputation methods. However, it is important to note that this finding is based on a limited set of six cross-modality imputation methods available in this benchmark. A recent study^51^ focused on cross-modality imputation has conducted a more extensive benchmark, evaluating 14 cross-modality prediction methods. This can be considered a complement to our study, although the SOTA method MIDAS from our study was not included in that analysis.

Future updating of such benchmarking including: (1) the impact of input dimensionality was not fully assessed in the current study, which are waiting to be carefully investigated in the future. In the scalability evaluation of the unpaired scRNA and scATAC mosaic integration, MIDAS was able to handle 10,000 paired and 10,000 unpaired cells in the BMMC Multiome simulation dataset, but it was limited in its availability for the SHARE-seq simulation dataset due to the higher dimensionality of the scATAC modality in the accuracy task than in the scalability task. This difference in dimensionality led to a significant increase in computational resource consumption for MIDAS. Future studies should more rigorously assess how input dimensionality affects algorithm performance. (2) Owing to limitations of available algorithms and the scarcity of publicly available high-quality spatial multiomics datasets, the current benchmark was limited in its evaluation of spatial multimodal integration methods, as well as comparison to non-spatial multimodal integration methods. We anticipate that forthcoming research will develop more efficient tools to handle spatial multi-omics data, and then a more comprehensive benchmark analyses for spatial multiomics integration can be conducted by incorporating unbiased and well-curated spatial multiomics datasets. Additional, for emerging single-cell multi-omics technologies and datasets, future benchmark studies should extend the scope to wider range of multi-omic data types, such as RNA and epigenetics data.

We recommend selecting optimal integration methods on the basis of their performance in usability, accuracy, and robustness (Extended Data Fig. 9). Computational hardware and resource requirements should also be considered, as the GPU devices and memory limits used in this study may not be applicable to all users.

In summary, our benchmark provides researchers with a comprehensive understanding of the available multimodal integration methods. We have developed a user-friendly web platform that allows researchers to access benchmark results and select the most appropriate integration method for their needs. Additionally, we have made available a reproducible pipeline for all benchmark methods and a Python package for evaluation metrics, enabling users to perform reproducible benchmarks on new datasets and integration algorithms.

### Protocol registration

The study was conducted according to the registered peer-reviewed protocol av ailable at https://springernature.figshare.com/articles/journal_contribution/Benchmar king_single-cell_multi-modal_data_integrations/26789572. Other than pre-registered and approved pilot data, all data reported in the paper were acquired after t he date of the registered protocol publication.

## Methods

### Preprocessing datasets

For multi-modal integration benchmark, we used nine publicly available single-cell multi-modal datasets for algorithms evaluation^41, 46, 52, 53, 54, 55^ (Supplementary Table 2, Supplementary Note 3). We tested different tasks for algorithms based on the modality type, cell annotation method, batch information, and dataset size for each benchmark dataset. Specifically, for biological conservation evaluation, if a benchmark dataset annotates cell types using joint embedding clusters or scRNA embedding clusters generated by single-cell analysis tools such as Seurat or Scanpy, the ground truth cell labels would favor the benchmark method using the same or similar cell clustering method, thus bias may occur. Therefore, we only used datasets with independent cell annotation methods as benchmark methods for biological conservation evaluation.

Our preprocessing of the multi-omics datasets followed the best practice of single-cell analysis across modalities^56^, as well as the user guidance provided in algorithm tutorials. For unpaired diagonal integration algorithms, which often involve transforming ATAC chromatin accessibility matrices into gene activity matrices (GAM), we computed the GAM using the GeneActivity function in Signac package^50^ using the ATAC per fragment information file. For datasets with raw SRA sequencing data only, we first applied NCBI SRA-toolkit to generate fastq files and cell ranger ARC pipeline (10x Genomics; v2.0.2) pipeline to generate the ATAC fragment information file. Then we computed the GAM using the GeneActivity function with the ATAC per fragment information file. For certain scATAC datasets with neither ATAC per fragment information file nor raw sequencing file, we used CreateGeneActivityMatrix function within the Seurat v3 package^37^ to compute GAM from the scATAC accessibility profile.

In the case of paired integration algorithms, typically reliant on top features, we utilized scanpy^57^ to calculate these features for Python-based algorithms, and employed Seurat v3^37^ for feature filtering in R-based algorithms.

### Usability assessment

To evaluate the usability of integration algorithms, we considered both user-friendliness metrics and scalability metrics for the algorithm. The user-friendliness metrics were adapted from previous benchmark study^58, 59^, including executability of the code (execute directly, executable with manual source code debugging or non-executable), quality of paper (peer review and performance benchmarking available), presence of tutorial for one or more examples to guide users and presence of function documentations for usage of all built-in functions.

The scalability metrics evaluate whether the method is executable for distinct dataset sizes (Supplementary Table 2) with limited computation hardware resource. The scalability of all multi-modal integration tools was assessed with CPU computation time, maximum memory use and GPU memory consumption. We used Linux GNU time command to collect the computation time and memory usage, and applied nvitop package in python to collect maximum GPU usage for all. For unpaired diagonal integration evaluation, we also evaluated the scalability for the GAM preparation step with GeneActivity^50^ and CreateGeneActivityMatrix^37^ function, as the generation of GAM also consumed long time and high memory for large scRNA and scATAC multimodal datasets. The scalability metrics evaluated the availability of all methods with the limits of 500GB RAM, 24 hours running time and 24 GB GPU memory.

Among single-cell multimodal integration tools, some algorithms can explain the relationship between input features from different modalities and each dimension of the cell latent space, which can be taken as a kind of biological interpretation. Similar to principal component analysis (PCA), each dimension can be “interpreted” by features with high absolute values in their weight, and features with no association with the dimension have weight values close to 0. In this study, we also assessed whether the benchmark algorithm is biologically interpretable in the output latent space, and report the interpretation method if the algorithm is interpretable. We scored algorithms with biological interpretability in latent space as 1, and introduce the specific method used for interpretation in the usability section. For algorithms with no biological interpretability in latent space, we scored 0 for this metric.

### Accuracy assessment

We employed both single embedding accuracy metrics and imputation accuracy (if imputation available) for paired multi-omics integration algorithms, employed both embedding accuracy metrics and cell alignment accuracy metrics for unpaired diagonal integration algorithms (Extended Data Fig. 10) and employed all metrics (if imputation available) for unpaired mosaic integration metrics. The particulars of these embedding accuracy metrics, cell alignment accuracy metrics and imputation accuracy metrics in this study are as follows:

#### Embedding accuracy metrics

For evaluating single embedding accuracy, we utilized two categories of metrics, focusing on batch effect removal and biological structure conversation^59^. Firstly, for batch effect removal evaluation, we employed cell-label-dependent metrics including batch ASW and KNN Graph Connectivity (GC), as well as cell-label-independent metric graph iLISI^59^. Also, for datasets with different time points or different disease states, we evaluated the batch effect removal metrics for samples of same time point or disease state, and then took the average from all time points or disease states as the output scores for the benchmark dataset. To reduce potential bias in cell labels and cell annotation methods, we used cell clusters to replace cell labels in each algorithm for batch ASW calculation. Secondly, for biological structure conversation evaluation, we applied global metrics as Adjusted Rand Index (ARI) and Adjusted Mutual Information (AMI), a local metric as graph cLISI, and a rare cell identity metric as isolated labels ASW. And these metrics were calculated for both original fine-grained cell annotations and broad cell types.

Among all embedding accuracy metrics, GC, graph iLISI and cLISI are computed from KNN (K-Nearest Neighbors) graphs rather than the latent embedding. The KNN graph is constructed by Euclidean distances on the joint embedding and is computed using the neighbor function. We used *k*=15 for graph connectivity and *k* as default value (1% of total cells) in graph_ilisi and graph_clisi function in scib package. Specifically, for paired integration methods, we calculated the biological conservation metrics both in the single batch dataset and mixed batches dataset to evaluate the performance of algorithms with and without batches. For algorithms correcting gene space and output graph rather than latent embedding, we took graph output of algorithm as input for all metrics evaluation except for ASW metrics.

*ASW*. The silhouette width quantifies the relative value of within-group distances of a cell and the between-group distances of that cell to the closest cluster. The silhouette width ranges from -1 to 1, with a score of 0 indicating that the group cannot be distinguished from other groups. The Average Silhouette Width (ASW) was computed as the average of the silhouette width for all cells.

For batch mixing evaluation, we initially adjusted the batch silhouette width of cell *i* by equation (1) and calculated the average batch silhouette width of all cells in cell cluster *j* as batch ASW_j_ (equation 2). The final batch ASW score (equation 3) was an average of batch ASW scores across all cell clusters. A higher batch ASW score indicates a lower remaining batch effect in the latent embedding.

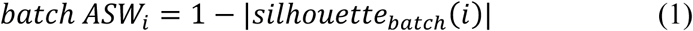

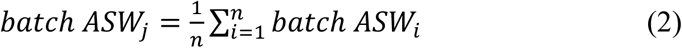

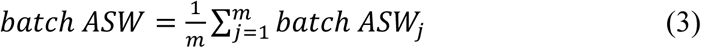

In this study, we considered the granularity of batch annotation for assessing batch effects, including batches from different samples and sites. For unpaired mosaic integration methods, we also used this metric to evaluate the batch removal effect between paired and unpaired multi-modal datasets, which treated each input dataset as a single batch.

To assess the methods’ ability to identify cell types in rare batches, we applied the isolated labels ASW metric. This metric grouped all cells as either isolated cells (representing cell types exhibited in the fewest batches) or non-isolated cells, and then calculate the ASW for the isolated cell group versus the non-isolated cell group. Then the ASW value was scaled to the range between 0 and 1 using the equation as *isolated labels ASW* = (*ASW* + 1)/2.

##### Graph connectivity

The KNN graph connectivity (GC) metric is used to evaluate batch effects through a KNN graph. For each cell type *c*, we constructed a subset KNN graph and calculated the percentage of the largest connected nodes in the graph. The GC was computed as following equation:

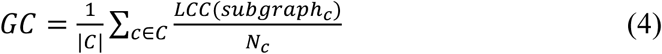

Here, *C* represents all cell types, *LCC*(*subgraph_c_*) represents largest connect components of the subgraph for the cell cluster *c*, and *N_c_* is the number of nodes in graph for cell type *c*. And we calculated the percentage of *LCC*(*subgraph_c_*) in *N_c_*. Then, GC was calculated as average percentages of connected cells in all cell types. This metric provides insight into the extent of batch effects captured by the KNN graph connectivity, where a higher GC score suggests a more connected graph and potentially lower batch effects. Similar with batch ASW metric, we used this metric to evaluate the unpaired mosaic integration. Also, we only applied this metric for datasets with well-curated and unbiased cell annotations, as this metric is influenced by the quality of cell type annotations.

##### Graph LISI

The Local Inverse Simpson’s Index (LISI) is computed from the neighborhood list in KNN graphs, which determines the number of cells selected from a local neighborhood list before a group can be observed twice. The LISI can be calculate as:

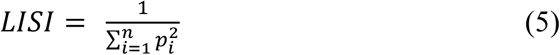

Where *n* is total number of groups, and *p_i_* is the proportion of cells from *i*th group in all cells. LISI ranges from 1 to *n* and larger LISI indicating better mixing of all groups. Adapted from scIB study, we set *n* as batches and normalize the LISI score using the following equation:

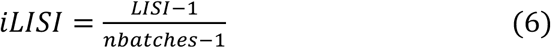

Here, iLISI represents the local batches mixture levels, where iLISI equals 1 when batches are mixed optimally. Furthermore, we set groups as cell types and scaled the LISI score to range from 0 to 1, defined as cLISI, to evaluate the biological structure conservation as following:

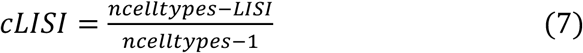

Here, cLISI equals 1 when LISI equals 1, indicating only single cell type clustering together locally. And cLISI equals to 0 when LISI equals to the number of all cell types, as all cell types mixed well in local KNN graphs.

##### ARI

The Adjusted Rand Index (ARI) compares Louvain cell clusters from each algorithm to the annotated cell labels. It considers both consistency of cell-cell pairs in the same cell clusters and consistency of cell-cell pairs in different cell clusters. An ARI of 0 indicates random cell clusters compared to annotated cell labels, while an ARI of 1 indicates a perfect match of cell clusters to cell labels. The ARI can be computed as the following equation:

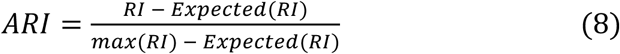

Here, *RI* represents the percentage of correct pairs for two cells in the same or different cell clusters in all possible cell-cell pairs. *Expected*(*RI*) represents *RI* score for randomly sampled cells, and *max*(*RI*) represents the percentage of correct pairs for a perfect match with cell labels.

##### AMI

Adjusted Mutual Information (AMI) also compares overlaps of cell clusters computed by each algorithm with annotated cell labels. It corrects mutual information by the mean of entropy from algorithm cell clusters and annotated cell labels using the following equation:

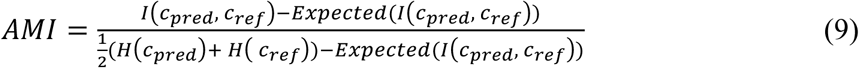

Here, *c_pred_* and *c_ref_* represent cell clusters predicted by each algorithm and reference cell labels. *i*(*c_pred_*, *c_ref_*) represents the mutual information of two cell groups, which is denoted as equation:

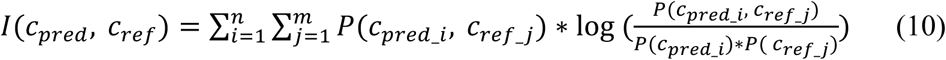

Where *i* and *j* indicate the *i*th cell cluster in *n* predicted cell groups the *j*th cell cluster in *m* reference cell groups. *P*(*c_pred_i_*, *c_ref_j_*) represents the percentage of cells in reference cell type *j* predicted as cell group *i* among all cells, while *P*(*c_pred_i_*) and *P*(*c_ref_j_*) represent the percentage of cells in predict cell group *i* and the percentage of cells in reference cell type *j* among all cells.

*H*(*c_pred_*) and *H*(*c_ref_*) in equation (9) represent the entropy of cell clusters in predictions and reference cell types. The equation of *H*(*c_pred_*) can be computed as follows:

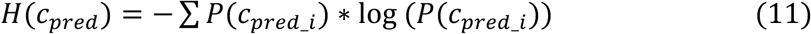

#### Cell alignment accuracy metrics

For unpaired multi-modal integration algorithms, we employed a combination of both embedding accuracy and alignment accuracy metrics. More precisely, the cell alignment accuracy metrics aim to evaluate the alignment accuracy of two modalities after integration^60^. These metrics evaluate the proportion of false cell match as Fraction of Samples Closer Than the True Match (FOSCTTM)^13^, and true match at the cell and cell type levels^60^. The details of cell alignment accuracy metrics are as follows:

##### FOSCTTM

FOSCTTM^61^ evaluates how closely the cell from different modalities are placed in the latent space by calculating the proportion of false match cells closer than the true match. Lower FOSCTTM score indicates a lower false match level and higher alignment accuracy, and a FOSCTTM of 0 indicates a perfect match between two modalities.

##### Cell match/Cell type match

For each cell in each modality, we first identified the nearest cell to a cell in the latent space, and determine whether the nearest cell in the other modality matches the cell type (cell type match) or cell barcode (cell match) of the select cell. As this metric is directional for integrated modalities, we calculated the true match proportion metric in each modality.

#### Imputation assessment

For paired integration, we simulated modality data sparsity by downsampling the sequencing counts in each cell to 10% of the original total counts, and then used benchmark integration methods to impute the downsampled modality. For unpaired mosaic integration, we constructed pseudo-unpaired multi-omics datasets from paired multi-omics data and impute different modalities from each modality of pseudo-unpaired datasets. For both paired integration methods and unpaired mosaic integration methods, we evaluated the accuracy of posterior imputation with gold standards of both raw data profile and KNN imputed profile from raw count data with the k-NN smoothing package^62^. For missing modality imputation, we also compared generative imputation results with KNN impute results in StabMap^42^, which imputed the missing modality from paired multi-modal data.

To provide a fair evaluation for imputation, we assessed the characteristics of both datasets and imputation methods. We calculated the nonzero ratios for raw and k-NN smoothed profile of scATAC modality for the benchmark datasets. If the nonzero ratio of scATAC profiles after k-NN smoothing is closer to 100%, indicating that most scATAC peaks in every cell are accessible, we did not use the k-NN smoothed profile of scATAC data as gold standards to avoid the overcorrection bias in the k-NN smoothed profiles. We also summarized the number of features imputed by each method, which reported the potential bias in imputation evaluation for method imputing too few features.

After above assessments, we applied different evaluation metrics for scRNA, scATAC, and ADT based on the nature of their modality. Specifically, for scRNA and ADT modalities that represent relative abundance, we calculated the Pearson correlation between raw expression and the algorithm’s posterior imputation. For the scATAC modality representing binary states of chromatin sites, we binarized the scATAC profile as binary gold standard, and then use an AUROC (area under receiver operating characteristic) metric to evaluation the accuracy of imputation. However, severe data sparsity may exist in the scATAC modality, and the positive chromatin accessible sites would be far less than negative inaccessible sites. For this condition, we adapted AUPR (area under precision recall) for imputation accuracy evaluation instead of the ATAC AUROC metric applied in the study of MIDAS paper^19^, which is less biased for unbalanced labels.

### Robustness assessment

We assessed the robustness of each method in two key aspects: (1) how consistent the method’s output is when run multiple times, and (2) how it performs under varying levels of data quality. The first aspect assessed algorithms of deep generative models with standard deviation (SD) of accuracy metrics. Non-deep algorithms often generate exactly same output when run multiple times, and the SD is equal to 0. For unpaired mosaic integration, we further evaluated the robustness of the methods across varying sizes of paired multi-modal datasets.

To simulate different levels of data sparsity in single cell multi-modal datasets, we first downsampled each modality to the sequencing depth of 75%, 50%, 25% and 10% compared to the original data. Subsequently, we evaluated the performance of algorithms under following three conditions: 1) when one modality is downsampled to a specific data sparsity, 2) when the other modality is downsampled to a specific data sparsity and 3) both modalities are downsampled simultaneously to a specific sparsity.

To quantitatively assess the robustness of each method across these multiple downsampling levels, we designed a multi-gradient AUC (Area Under Curve) metric. This metric was calculated by following two steps: 1) it computes the percentage of accuracy values for a selected accuracy metric (ARI for etc.) at distinct downsampling levels comparing to original sequencing depth level; 2) it computes the AUC score for the curve of selected accuracy metric under all downsampling levels. The maximum of percentages under distinct levels was scaled to 1 if the score of distinct downsampling levels higher than raw data. An AUC score of 1 signifies that the algorithm demonstrates complete robustness to data sparsity, maintaining the same or higher performance across all downsampling levels for the given metric.

### Metrics summary and ranking

In the usability evaluation, we assessed both user-friendliness and scalability metrics. User-friendliness metric scores were quantified using a binary (1 and 0) score for each metric, and the average of all metrics was calculated. If a method is non-executable, we provided detailed reasons and exclude it from all other evaluations. Scalability scores were quantified and ranked based on the number of available tasks for each method. The actual consumption of RAM memory, GPU memory, and running time were also reported in the Results section and supplementary tables of our manuscript.

In the accuracy evaluation, we reported the original accuracy metric scores for benchmark algorithms, and then ranked these algorithms by the accuracy metric scores of biological conservation, batch effect removal and cell alignment accuracy metrics in all benchmark datasets.

To get the accuracy metrics ranking from metrics of different ranges, we first applied min-max normalize to each metric for the *i*th benchmark algorithm as follows:

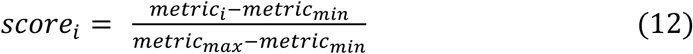

The algorithm’s score was normalized by the scores of the best and worst-performing algorithms in each metric, with the best method receiving a score of 1 and the worst a score of 0. Second, we calculated the average score for normalized biological conservation, batch effect removal and cell alignment accuracy metrics separately in all tested datasets for ranking. For paired integration accuracy, we employed the biological conservation score calculated from both single-batch and mixed batches datasets. Finally, we reported the original metric scores and average rankings of biological conservation, batch effect removal and cell alignment accuracy for each method respectively.

In the robustness evaluation, we applied the same equations as accuracy evaluation section to compute the accuracy metrics and scores for the downsampled datasets. However, paired multi-modal integration evaluation only used the appropriate metrics filtered by the methods mentioned in the *robustness assessment section*. The final robustness score was computed from the accuracy score at the lowest sequencing depth (*S*_*scc*_10_), which evaluates the method’s accuracy under the worst data quality conditions, and the average multi-gradient AUC score from selected metrics at all downsampled sequencing depths (*S_grad_auc_*). We reported the robustness score (*S_grad_auc_* and *S_acc_10_*) and ranks of selected metrics for each benchmark method, and ranked robustness of benchmark methods by the average ranks of two robustness score as follows:

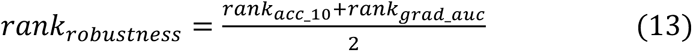

For unpaired mosaic integration, we further assessed the robustness of tested methods to paired multi-modal dataset size using the average multi-gradient AUC score (*S_size_auc_*). And robustness of unpaired mosaic integration methods was finally ranked by the average ranks of three robustness score as follows:

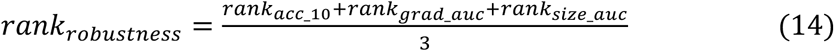

### Computational resource

Most of the benchmark tasks were conducted on an Ubuntu18 system of with 2×52 cores CPU, 1000G RAM and 4 NVIDIA 3090 24G GPU. To ensure that these algorithms can be applied in common scenarios, we set a maximum computational resource limit of 500GB RAM and a single NVIDIA 3090 24GB GPU for each algorithm. Some algorithms that only support python2 and older CUDA versions were run on a server with 2 NVIDIA 1080Ti 12GB GPU, 2×52 cores and 500 G RAM. Both servers operated on the Ubuntu 18.04 operating system.

## Data availability

BMMC 10X Multiome and CITE-seq datasets^52^ analyzed in this study are available at https://openproblems.bio/events/2021-09_neurips/. The raw sequencing files of BMMC Multiome datasets used in this study are available at GEO database with accession number GSE194122. The HSPC 10X Multiome and CITE-seq datasets^53^ are available at https://www.kaggle.com/competitions/open-problems-multimodal/data. The SHARE-seq skin data^54^ can be downloaded from GEO database with accession number GSM4156608 and GSM4156597. The COVID19 CITE-seq data^55^ is available at E-MTAB-10026 (ArrayExpress). The human WBC CITE-seq data^38^ is available at https://atlas.fredhutch.org/nygc/multimodal-pbmc/. The 10X NSCLC CITE-seq, 10X kidney cancer CITE-seq, 10X mouse brain Multiome and 10X PBMC Multiome datasets are downloaded from 10X Genomics website (https://www.10xgenomics.com/datasets/). For spatial multi-omic integration tasks, we obtained SPOTS mouse spleen data^46^ from GSE198353(GEO), mouse thymus data^41^ from https://zenodo.org/records/10362607, and human lymph node data^41^ from https://drive.google.com/drive/folders/1RlU3JmHg_LZM1d-o6QORvykYPoulWWMI. The processed input datasets for all benchmark methods are available at a publicly available Figshare repository (https://figshare.com/projects/Single-cell_multimodal_integration_benchmark_SCMMIB_register_report_Stage_2_study_/221476).

## Code availability

We have uploaded the source code for the evaluation metrics Python package and the scripts for reproducing figures in the Stage 2 manuscript to a GitHub repository at https://github.com/bm2-lab/SCMMI_Benchmark/. Additionally, a pipeline for running all benchmark methods has been uploaded to https://github.com/bm2-lab/SCMMIB_pipeline. The interactive website for detailed supplementary results of this study is available at https://bm2-lab.github.io/SCMMIB-reproducibility/. Code is also available in the Zenodo repository via https://doi.org/10.5281/zenodo.1479295163.

## Acknowledgements

1. Q. L. was supported by National Natural Science Foundation of China (Grant No. T24250193, 32341008), the National Key Research and Development Program of China (Grant No. 2021YFF1201200, No. 2021YFF1200900), Shanghai Pilot Program for Basic Research, Shanghai Science and Technology Innovation Action Plan-Key Specialization in Computational Biology, Shanghai Shuguang Scholars Project, Shanghai Excellent Academic Leader Project, Shanghai Municipal Science and Technology Major Project (Grant No. 2021SHZDZX0100) and Fundamental Research Funds for the Central Universities. S. F. was supported by National Natural Science Foundation of China (Grant No. 32400521) and China Postdoctoral Science Foundation (Grant No. 2023M742651, GZC20231946).

## Author contributions

S.F., S.W. and Q.L. conceived the project. S.F., S.W. designed the Stage 1 proposal and performed Stage 2 data analysis with help from D.S., G.L. and Y.G. S.F., S.W. and Q.L. wrote the manuscript with input from all authors. Q.L. supervised the entire project. All authors read and approved the final manuscript.

