## Extended and supplementary figures for "Benchmarking single-cell multi-modal data integrations": Extend_data_table_and_figures.pdf

### Extended data figures

| Dataset name | Number of cells | Number of features | | | Non-zero value ratio | | |
|---|---|---|---|---|---|---|---|
| | | scRNA | scATAC | GAM | scRNA | scATAC | GAM |
| BMMC Multiome p10 | 6,924 | 13,431 | 116,490 | 19,607 | 0.082 | 0.031 | 0.256 |
| BMMC Multiome s3d10 | 6,781 | 13,431 | 116,490 | - | 0.102 | 0.032 | - |
| HSPC p10 Multiome | 10,587 | 23,418 | 228,942 | 25,411 | 0.165 | 0.025 | 0.156 |
| SHARE-seq skin | 34,774 | 23,296 | 344,592 | 22,499 | 0.028 | 0.012 | 0.099 |
| 10X Mouse Brain | 12,138 | 32,286 | 66,909 | 21,807 | 0.088 | 0.024 | 0.154 |
| 10X PBMC | 15,021 | 36,601 | 115,980 | 19,607 | 0.05 | 0.054 | 0.298 |
| BMMC Multiome p10 10% | 6,924 | 13,431 | 116,490 | 21,372 | 0.012 | 0.006 | 0.029 |
| BMMC Multiome p10 25% | 6,924 | 13,431 | 116,490 | 21,372 | 0.027 | 0.014 | 0.062 |
| BMMC Multiome p10 50% | 6,924 | 13,431 | 116,490 | 21,372 | 0.048 | 0.023 | 0.099 |
| BMMC Multiome p10 75% | 6,924 | 13,431 | 116,490 | 21,372 | 0.066 | 0.028 | 0.12 |

**Extended Data Fig. 1 | Summary table of scRNA and scATAC multimodal datasets analyzed in the SCMMIB study.** Details of cell numbers, feature counts and non-zero ratios in all scRNA and scATAC datasets for paired or unpaired integration methods evaluations. Datasets used in unpaired scRNA and scATAC diagonal integration were further evaluated using the attributes of gene activity matrix (GAM).

| Dataset name | Number of cells | Number of features | | Non-zero value ratio | |
|---|---|---|---|---|---|
| | | scRNA | ADT | scRNA | ADT |
| BMMC CITE-seq p10 | 9,026 | 13,953 | 134 | 0.104 | 0.839 |
| BMMC CITE-seq s2d1 | 10,465 | 13,953 | 134 | 0.085 | 0.71 |
| HSPC p10 CITE-seq | 7,099 | 22,002 | 140 | 0.22 | 0.832 |
| Human WBC | 161,764 | 20,729 | 228 | 0.1 | 0.911 |
| 10X NSCLC | 15,618 | 36,601 | 9 | 0.096 | 0.946 |
| COVID19 CITE-seq | 647,366 | 24,737 | 192 | 0.053 | 0.881 |
| 10X kidney cancer | 20,974 | 37,143 | 32 | 0.14 | 0.881 |
| Lymph node spatial | 6,843 | 18,085 | 29 | 0.085 | 0.977 |
| Spleen SPOTS | 5,336 | 32,285 | 10 | 0.105 | 0.999 |
| Thymus spatial | 17,824 | 26,857 | 30 | 0.033 | 0.298 |
| BMMC CITE-seq p10 10% | 9,026 | 13,953 | 134 | 0.02 | 0.416 |
| BMMC CITE-seq p10 25% | 9,026 | 13,953 | 134 | 0.039 | 0.599 |
| BMMC CITE-seq p10 50% | 9,026 | 13,953 | 134 | 0.065 | 0.732 |
| BMMC CITE-seq p10 75% | 9,026 | 13,953 | 134 | 0.086 | 0.798 |

**Extended Data Fig. 2 | Summary table of scRNA and ADT multimodal datasets analyzed in the SCMMIB study.** Details of cell numbers, feature counts and non-zero ratios in all scRNA and ADT datasets for paired scRNA and ADT integration benchmark.

| Dataset name | Number of cells (pair data, unpair data) | Number of features | | Non-zero value ratio | |
|---|---|---|---|---|---|
| | | scRNA | 2nd modality (scATAC/ADT) | scRNA | 2nd modality (pair data, unpair data) |
| BMMC-CITE-s2d1_s3d6 | 11,035, 10,465 | 13,953 | 134 | 0.094,0.085 | 0.784,0.71 |
| BMMC-Multiome-s1d1_s3d10 | 6224, 6781 | 13,431 | 116490 | 0.102,0.102 | 0.028,0.032 |
| SHARE-Multiome-c5k_cnk | 3000-20,000,5000 | 23,296 | 344,592 | 0.028,0.027 | 0.012,0.012 |
| BMMC-Multiome-c5k_cnk | 3000-20,000,5000 | 13,431 | 116,490 | 0.081,0.083 | 0.031,0.031 |
| BMMC-CITE_seq-c5k_cnk | 3000-20,000,5000 | 13,953 | 134 | 0.105,0.104 | 0.841,0.841 |
| BMMC-Multiome s1d1_s3d10 10% | 6,224, 6,781 | 13,431 | 116,490 | 0.016,0.015 | 0.006,0.006 |
| BMMC-Multiome s1d1_s3d10 25% | 6,224, 6,781 | 13,431 | 116,490 | 0.035,0.035 | 0.013,0.013 |
| BMMC-Multiome s1d1_s3d10 50% | 6,224, 6,781 | 13,431 | 116,490 | 0.062,0.061 | 0.021,0.023 |
| BMMC-Multiome s1d1_s3d10 75% | 6,224, 6,781 | 13,431 | 116,490 | 0.083,0.083 | 0.026,0.029 |
| BMMC CITE-seq s2d1_s3d6 10% | 11,035,10,465 | 13,953 | 134 | 0.017,0.016 | 0.272,0.262 |
| BMMC CITE-seq s2d1_s3d6 25% | 11,035,10,465 | 13,953 | 134 | 0.035,0.031 | 0.475,0.43 |
| BMMC CITE-seq s2d1_s3d6 50% | 11,035,10,465 | 13,953 | 134 | 0.058,0.052 | 0.641,0.574 |
| BMMC CITE-seq s2d1_s3d6 75% | 11,035,10,465 | 13,953 | 134 | 0.078,0.069 | 0.729,0.655 |
| SHARE-Multiome-c5k_c5k 10% | 5,000, 5,000 | 23,296 | 344,592 | 0.004,0.004 | 0.001,0.001 |
| SHARE-Multiome-c5k_c5k 25% | 5,000, 5,000 | 23,296 | 344,592 | 0.008,0.008 | 0.003,0.003 |
| SHARE-Multiome-c5k_c5k 50% | 5,000, 5,000 | 23,296 | 344,592 | 0.015,0.015 | 0.006,0.006 |
| SHARE-Multiome-c5k_c5k 75% | 5,000, 5,000 | 23,296 | 344,592 | 0.022,0.022 | 0.009,0.009 |

**Extended Data Fig. 3 | Summary table of multimodal datasets analyzed in mosaic multimodal integration tasks in the SCMMIB study.** Details of cell numbers, feature counts and non-zero ratios of datasets for unpaired mosaic integration benchmark. The characteristics of paired and unpaired simulation datasets were separated with comma.

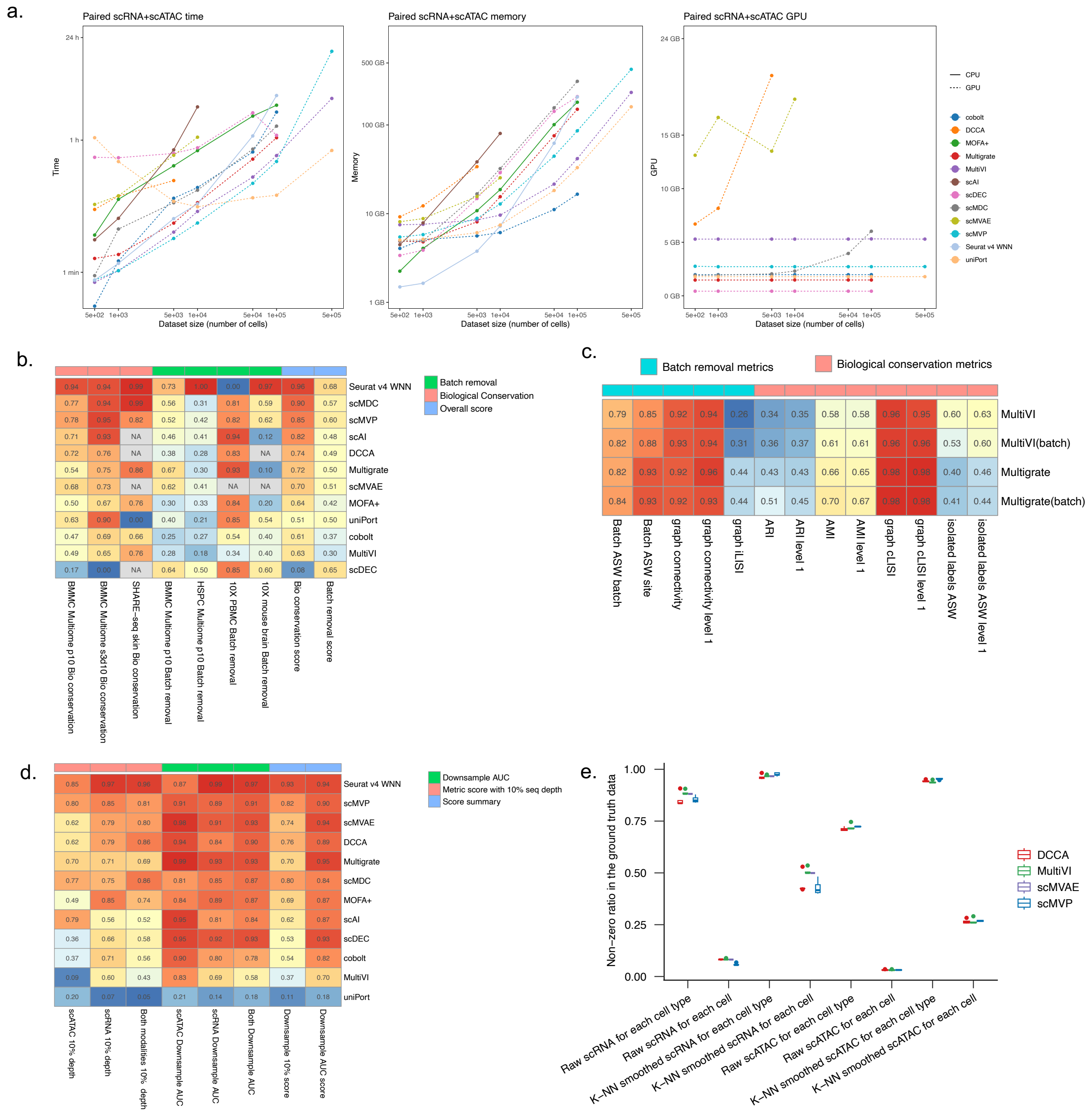


**Extended Data Fig. 4 | Details of scalability, accuracy, and robustness metrics for the paired scRNA and scATAC integration methods.** (**a**) Scalability line plot of running time, peak memory, and GPU memory for all paired scRNA and scATAC integration algorithms. Algorithms using GPU acceleration are plotted with dashed lines. (**b**) Heatmap of summarized accuracy metrics for paired scRNA and scATAC integration methods in Fig. 2b. The summarized metric scores are shown in each cell of the heatmap. (**c**) Evaluation for algorithms with optional batch parameters. The metric results calculated with batch information input are labeled with the "(batch)" suffix in figure. (**d**) Heatmap of summarized robustness metrics for paired scRNA and scATAC integration methods in Fig. 2c. The summarized metric scores are shown in each cell of the heatmap. (**e**) Non-zero ratios of the ground truth data used for paired scRNA and scATAC imputation evaluation.

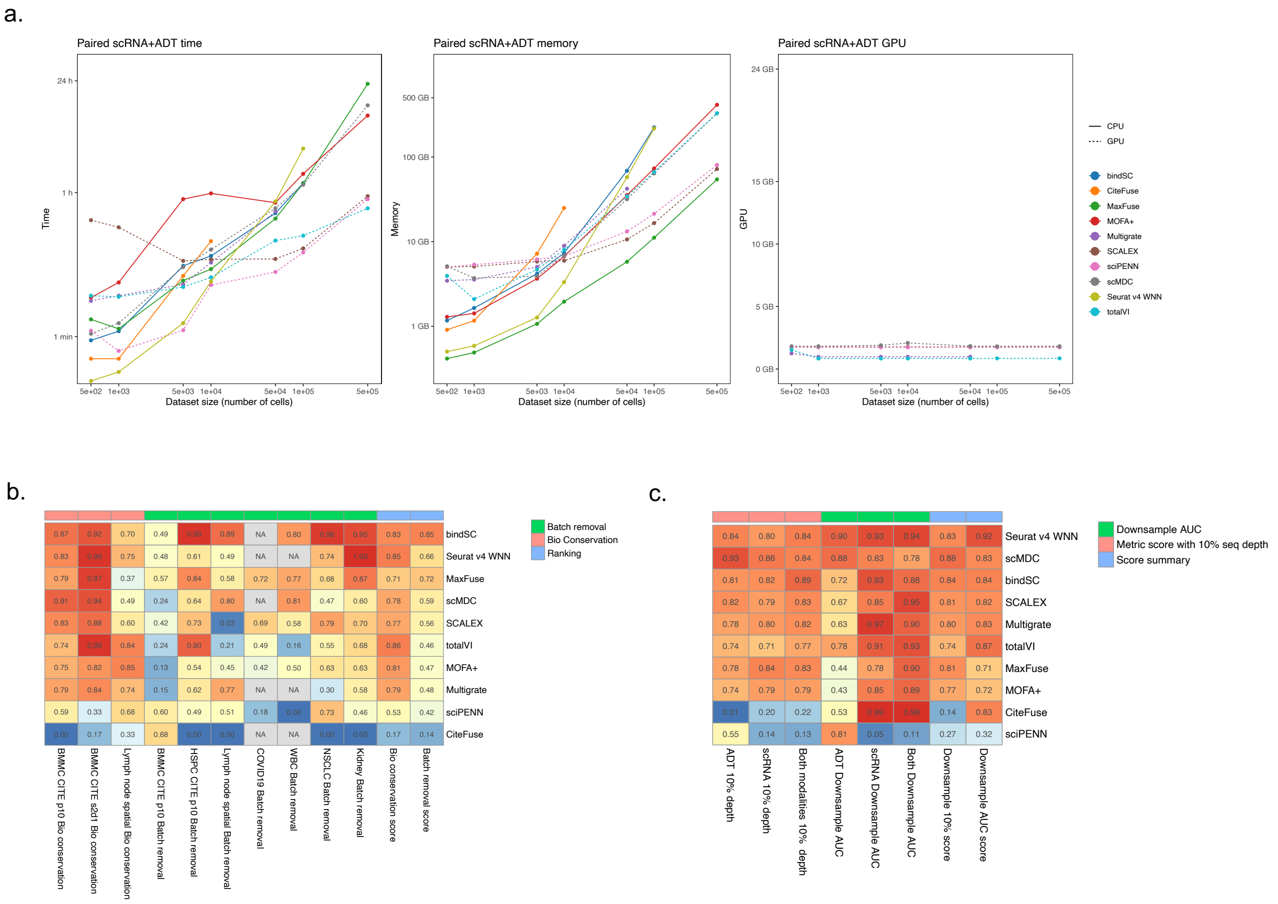


**Extended Data Fig. 5 | Details of scalability, accuracy, and robustness metrics for the paired scRNA and ADT integration methods.** (**a**) Scalability line plot of running time, peak memory, and GPU memory for all paired scRNA and ADT integration algorithms. Algorithms using GPU acceleration are plotted with dashed lines. (**b**) Heatmap of summarized accuracy metrics for paired scRNA and ADT integration methods in Fig. 3b. The summarized metric scores are shown in each cell of the heatmap. (**c**) Heatmap of summarized robustness metrics for paired scRNA and ADT integration methods in Fig. 3d.

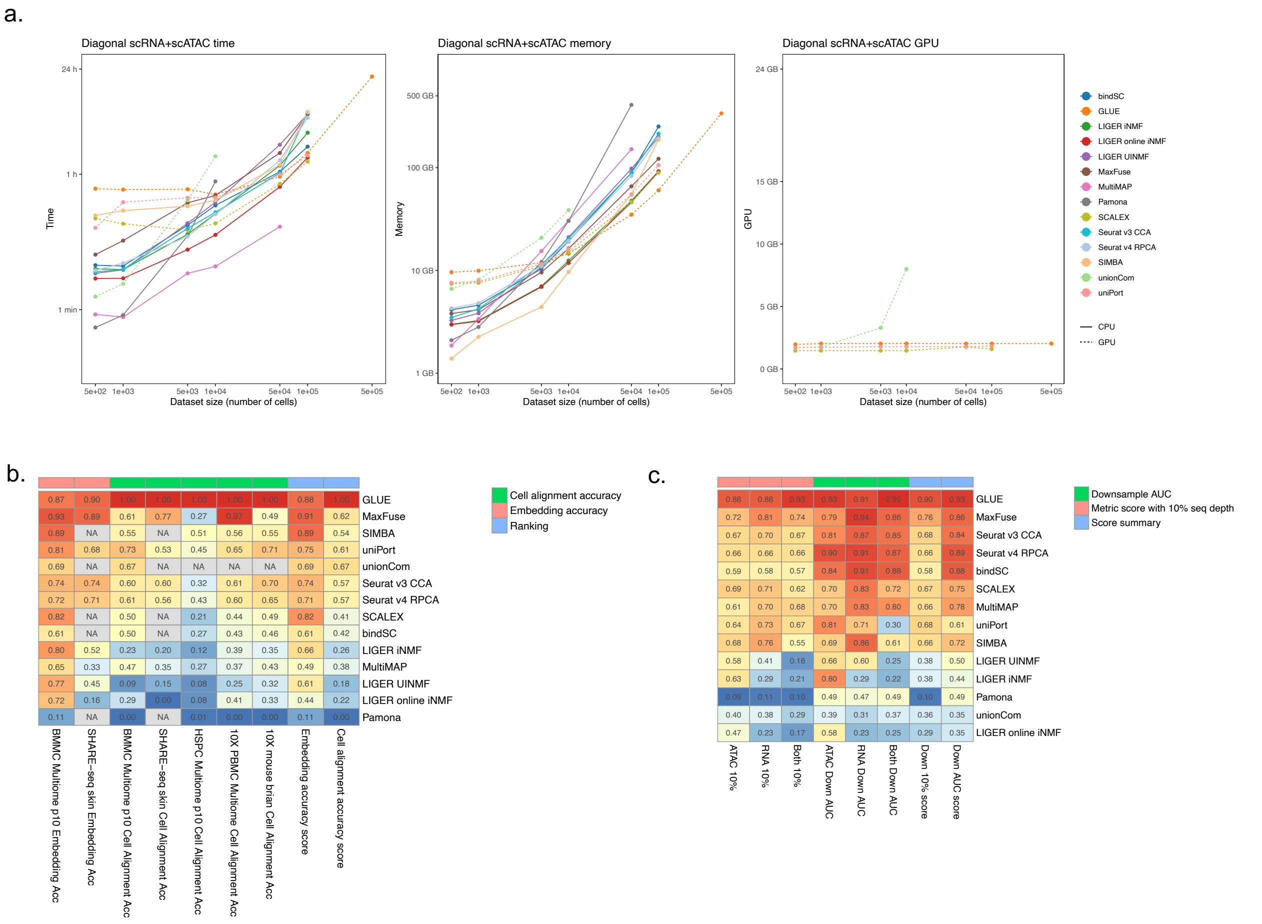


**Extended Data Fig. 6 | Details of scalability, accuracy, and robustness metrics for unpaired scRNA and scATAC diagonal integration methods.** (**a**) Scalability line plot of running time, peak memory, and GPU memory for all unpaired scRNA and scATAC diagonal integration algorithms. Algorithms using GPU acceleration are plotted with dashed lines. (**b**) Heatmap of summarized accuracy metrics for unpaired scRNA and scATAC diagonal integration methods in Fig. 4c. The summarized metric scores are shown in each cell of the heatmap. (**c**) Heatmap of summarized robustness metrics for unpaired scRNA and scATAC diagonal integration methods in Fig. 4f.

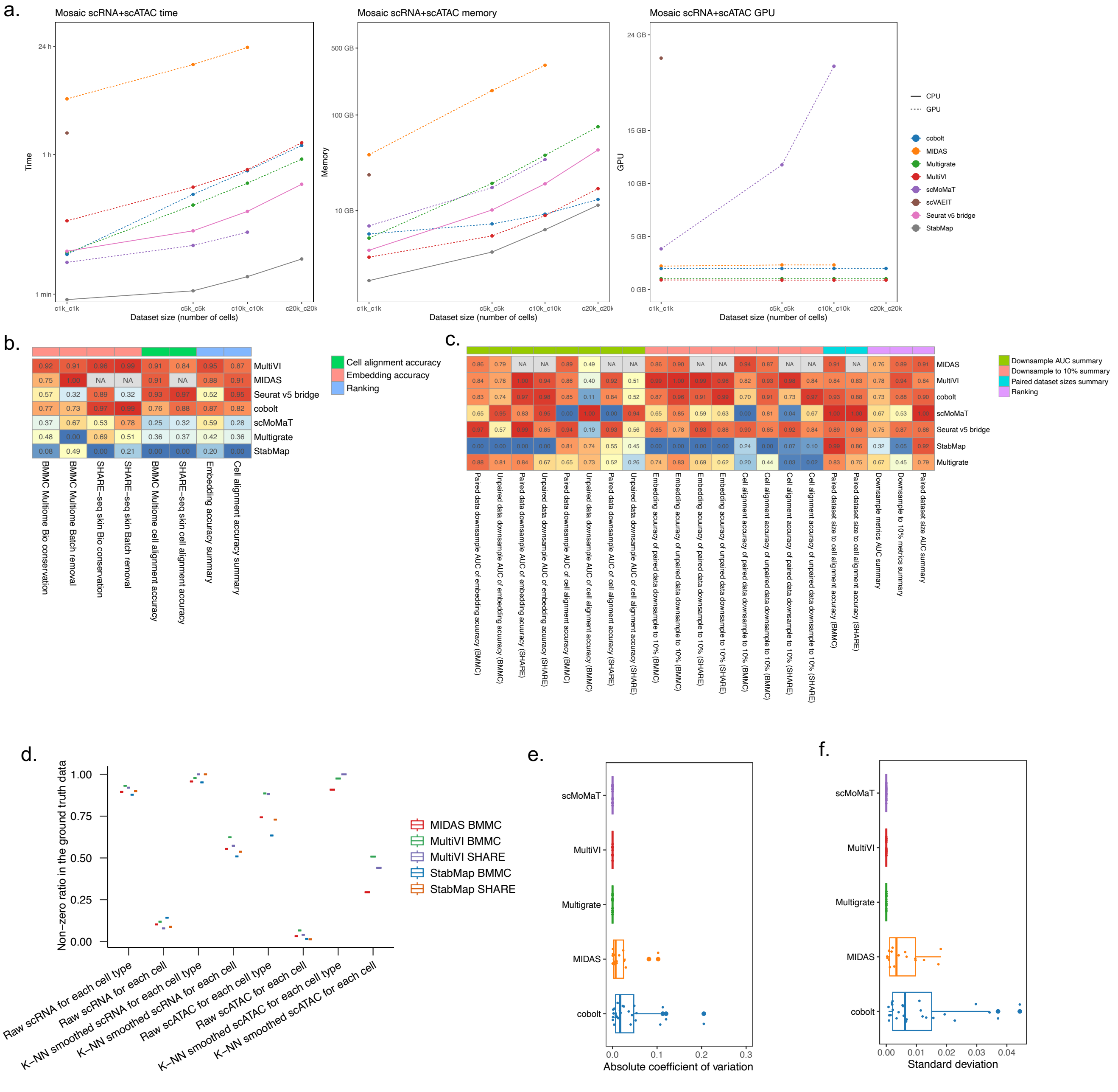


**Extended Data Fig. 7 | Details of usability, accuracy, and robustness metrics for the unpaired scRNA and scATAC mosaic integration methods.** (**a**) Scalability line plot of running time, peak memory, and GPU memory for all unpaired scRNA and scATAC mosaic integration algorithms. Algorithms using GPU acceleration are plotted with dashed lines. (**b**) Heatmap of summarized accuracy metrics for unpaired scRNA and scATAC mosaic integration methods in Fig. 5b. The summarized metric scores are shown in each cell of the heatmap. (**c**) Heatmap of summarized robustness metrics for unpaired scRNA and scATAC mosaic integration methods in Fig. 5e. The summarized metric scores are shown in each cell of the heatmap. (**d**) Non-zero ratios of the ground truth data used for mosaic scRNA and scATAC imputation evaluation. **e-f.** Stability of algorithm embedding output in 5 repeated runs, evaluated with (**e**) absolute coefficient of variation and (**f**) standard deviation of metric values for all accuracy metrics.

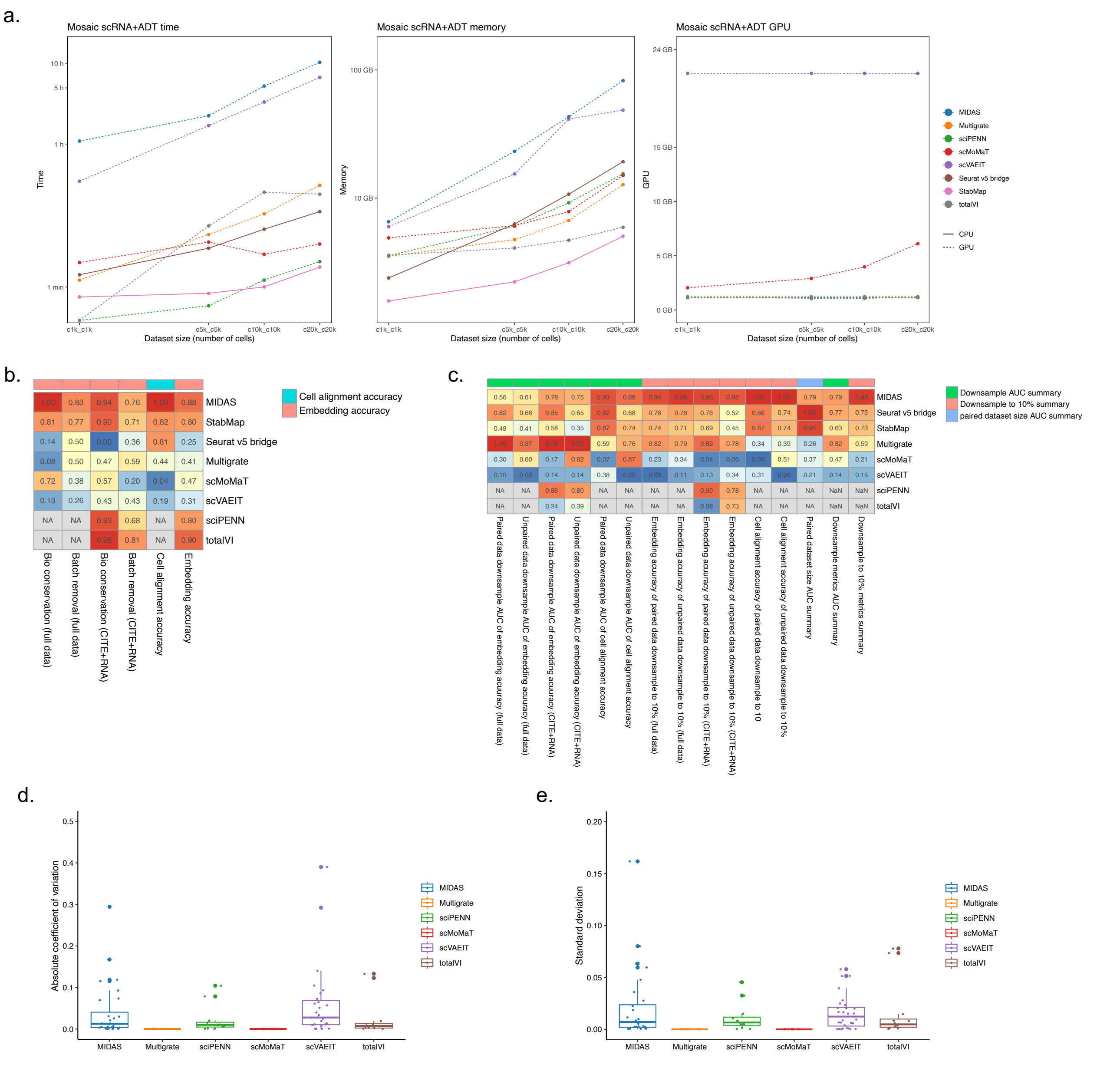


**Extended Data Fig. 8 | Details of usability, accuracy and robustness metrics for the unpaired scRNA and ADT mosaic integration methods.** (**a**) Scalability line plot of running time, peak memory, and GPU memory for all unpaired scRNA and ADT mosaic integration algorithms. Algorithms using GPU acceleration are plotted with dashed lines. (**b**) Heatmap of summarized accuracy metrics for unpaired scRNA and ADT mosaic integration methods in Fig. 6b. The summarized metric scores are shown in each cell of the heatmap. (**c**) Heatmap of summarized robustness metrics for unpaired scRNA and ADT mosaic integration methods in Fig. 6c. The summarized metric scores are shown in each cell of the heatmap. **d-e.** Stability of algorithm embedding output in 5 repeated runs, evaluated with (**d**) absolute coefficient of variation and (**e**) standard deviation of metric values for all accuracy metrics.

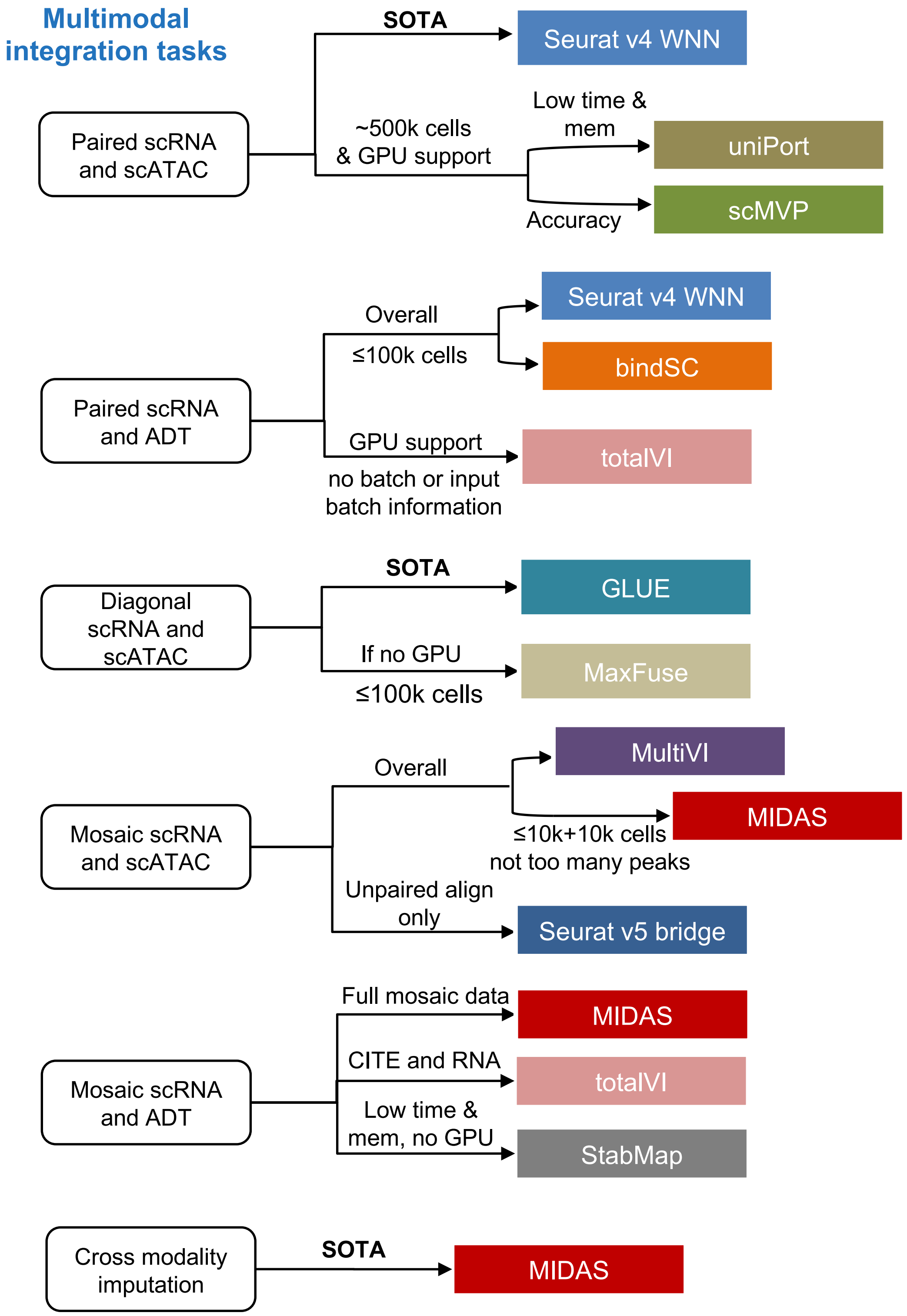


**Extended Data Fig. 9 | Guidelines for single-cell multimodal integrations.** Recommendations for the optimal method in certain integration task. The methods were recommended based on overall rankings in usability, accuracy, and robustness. For more specific details, users should refer to the conclusions in the corresponding results section. SOTA: state-of-the-art method.

| Metric | Group | Characteristic | Input format |
| --- | --- | --- | --- |
| batch ASW | batch effect removal | cell-cluster-dependent | Latent embedding |
| GC | batch effect removal | cell-label-dependent | KNN graph |
| graph iLISI | batch effect removal | local metric | KNN graph |
| ARI | biological conservation | global metric | Latent embedding |
| AMI | biological conservation | global metric | Latent embedding |
| graph cLISI | biological conservation | local metric | KNN graph |
| isolated labels ASW | biological conservation | rare cell type | Latent embedding |
| FOSCTTM | cell alignment accuracy | proportion of false match | Latent embedding |
| cell match/cell type match | cell alignment accuracy | proportion of true match | Latent embedding |
| AUPR/AUROC | imputation accuracy | accuracy of peaks' ranking | ATAC profile |
| Pearson's r | imputation accuracy | correlation to ground truth | RNA or ADT profile |

**Extended Data Fig. 10 | Summary table of accuracy metrics used in this study.**
