## Extended and supplementary figures for "Benchmarking single-cell multi-modal data integrations": Supplementary Figures.pdf

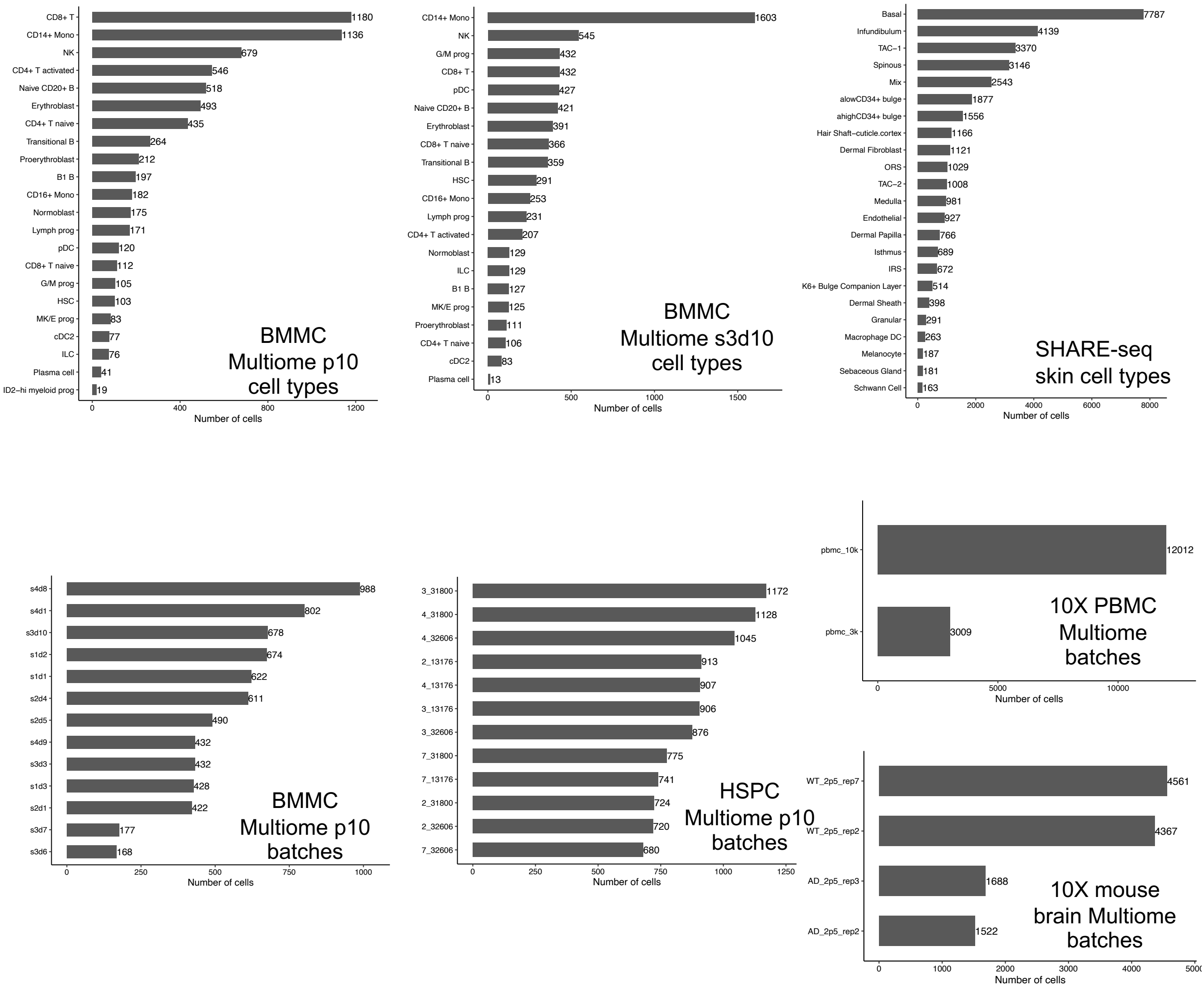


Supplementary Fig. 1. Summary of cell number in batches and cell types of scRNA and scATAC multimodal datasets used in the SCMMIB study. The y axis of boxplot shows the name of batch or cell type, and the x axis represented the number of cells.

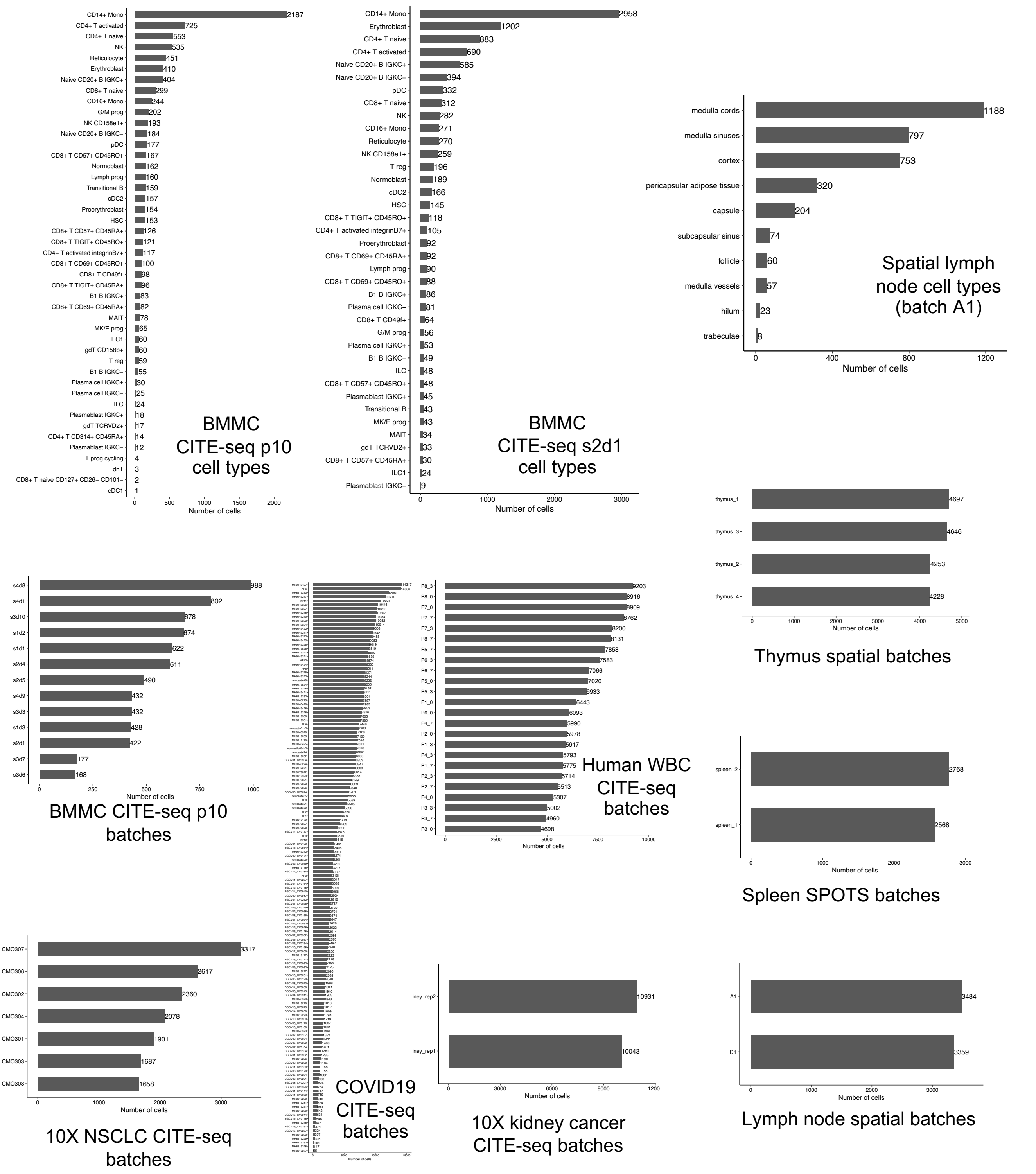


Supplementary Fig. 2. Summary of cell number in batches and cell types of scRNA and ADT datasets as well as spatial scRNA and ADT datasets used in the SCMMIB study. The y axis of boxplot shows the name of batch or cell type, and the x axis represented the number of cells.

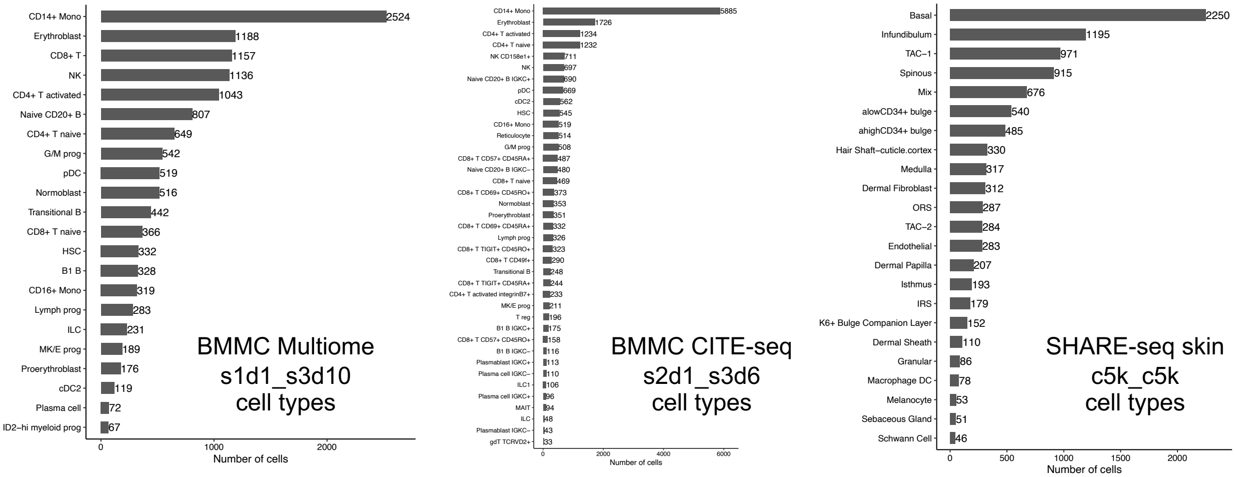


Supplementary Fig. 3. Summary of cell number in cell types of mosaic integration simulation datasets used in the SCMMIB study. The y axis of boxplot shows the name of cell type, and the x axis represented the number of cells.

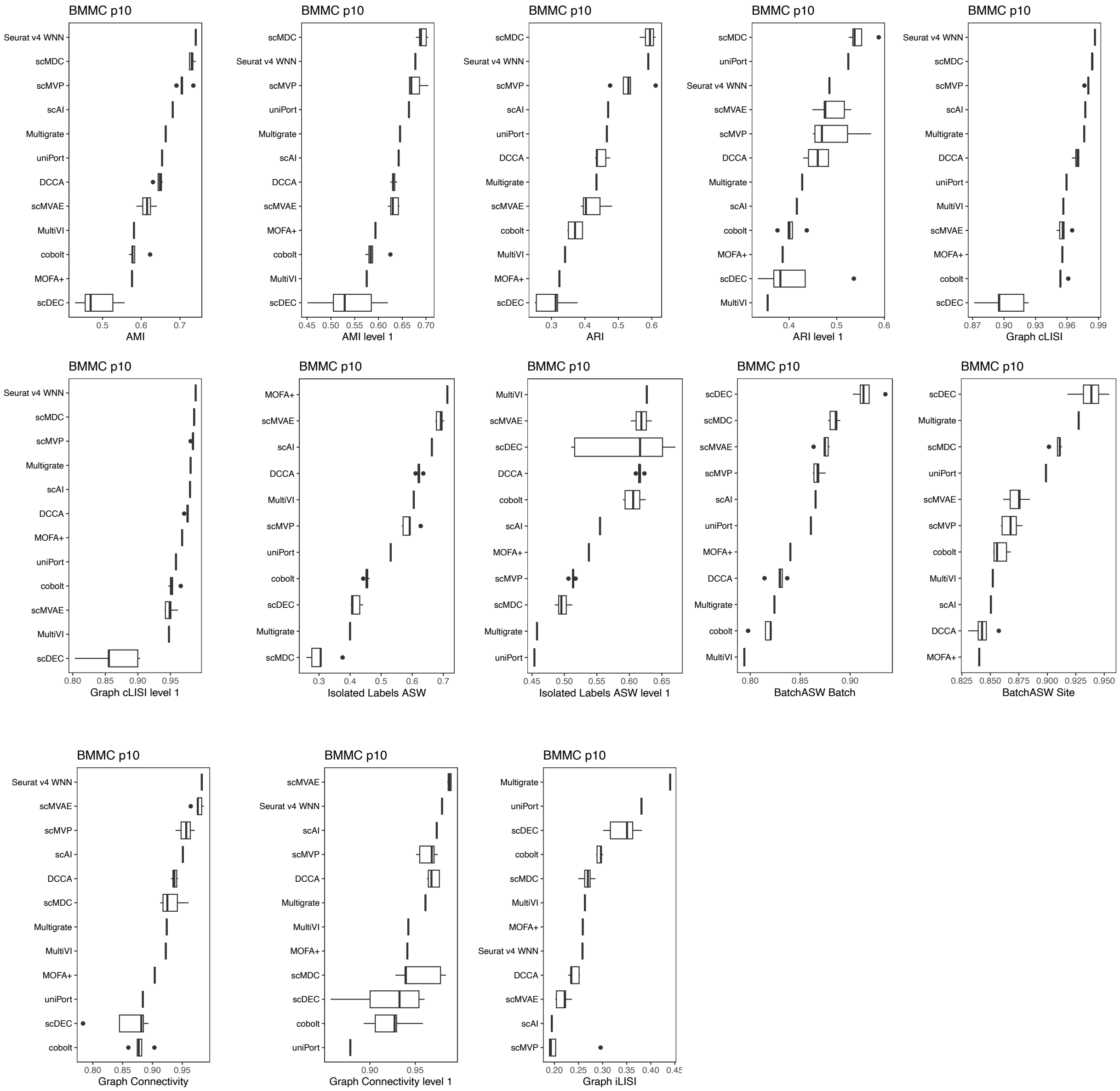


Supplementary Fig. 4. Boxplot of accuracy metrics for paired scRNA and scATAC integration algorithms in BMMC Multiome p10 dataset, corresponding to Fig. 2b and Extended Data Fig. 4b,

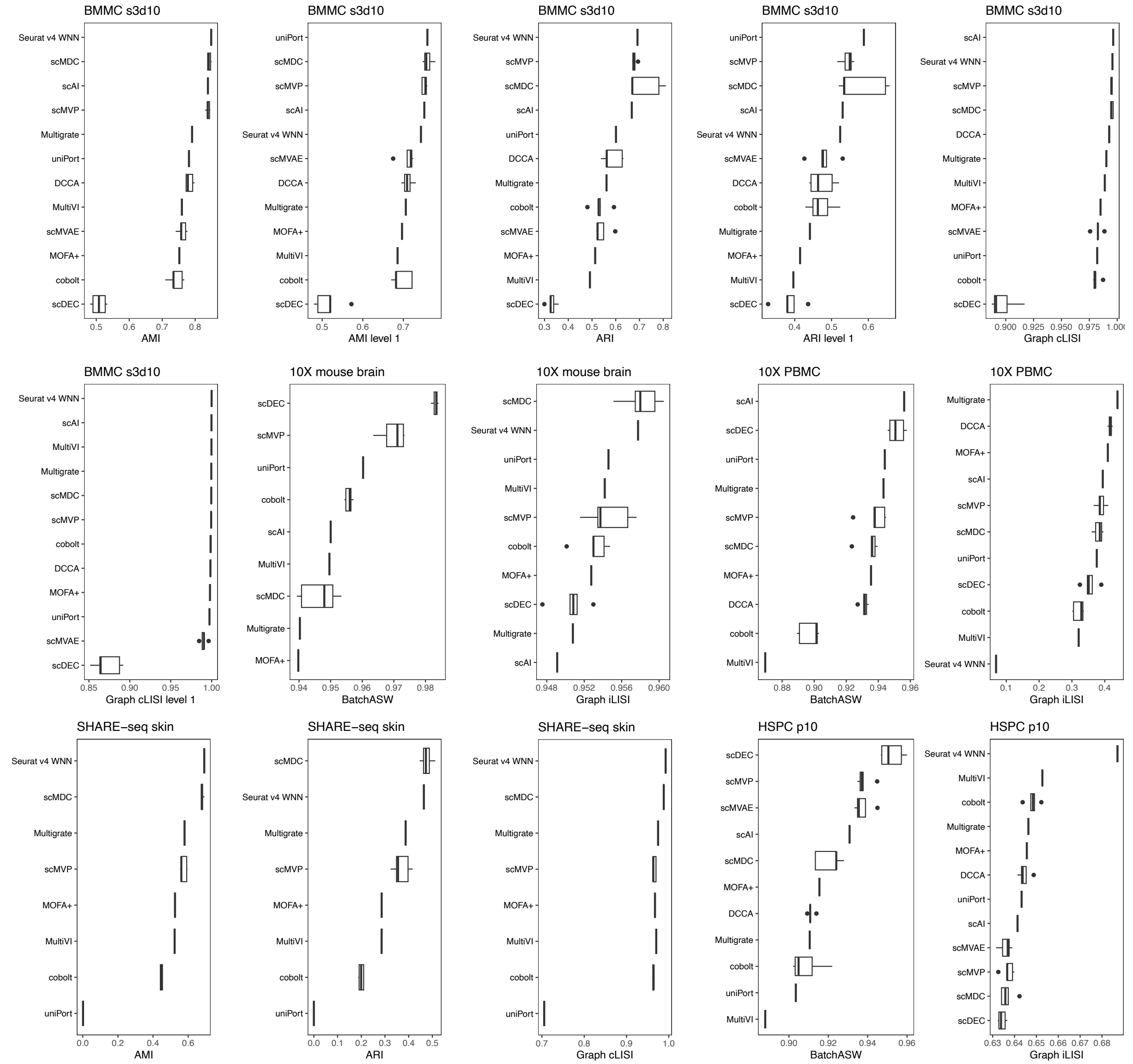


Supplementary Fig. 5. Boxplot of accuracy metrics for paired scRNA and scATAC integration algorithms in other datasets except for BMMC Multiome p10, corresponding to Fig. 2b and Extended Data Fig. 4b. The dataset names are labelled at the top of each boxplot.

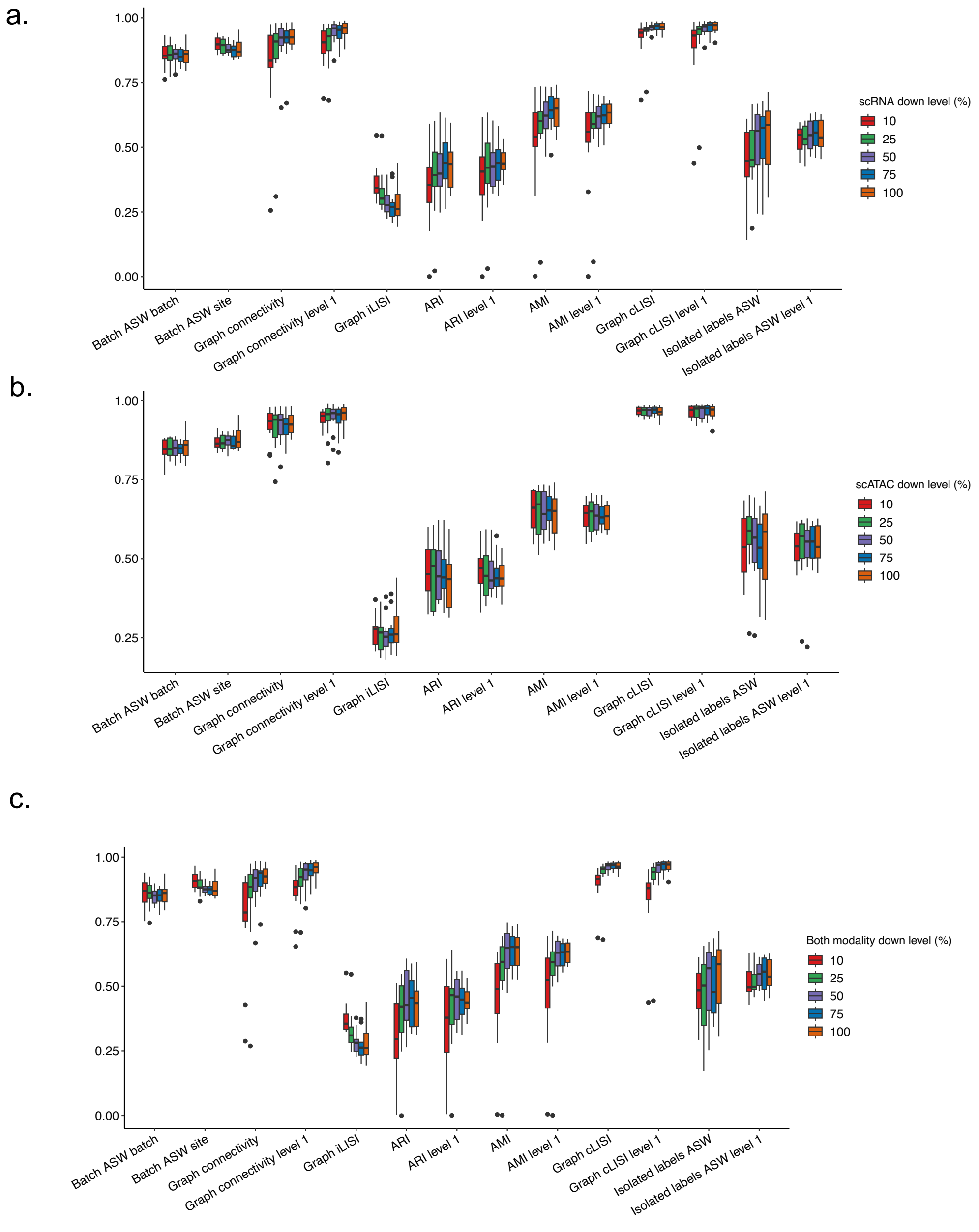


Supplementary Fig. 6. Boxplot of the biological conservation and batch removal metric values for all paired scRNA and scATAC integration methods at all downsampling levels. This figure shows the metric scores distribution for simulations of **(a)** scRNA modality downsampling, **(b)** scATAC modality downsampling and **(c)** both modalities downsampling.

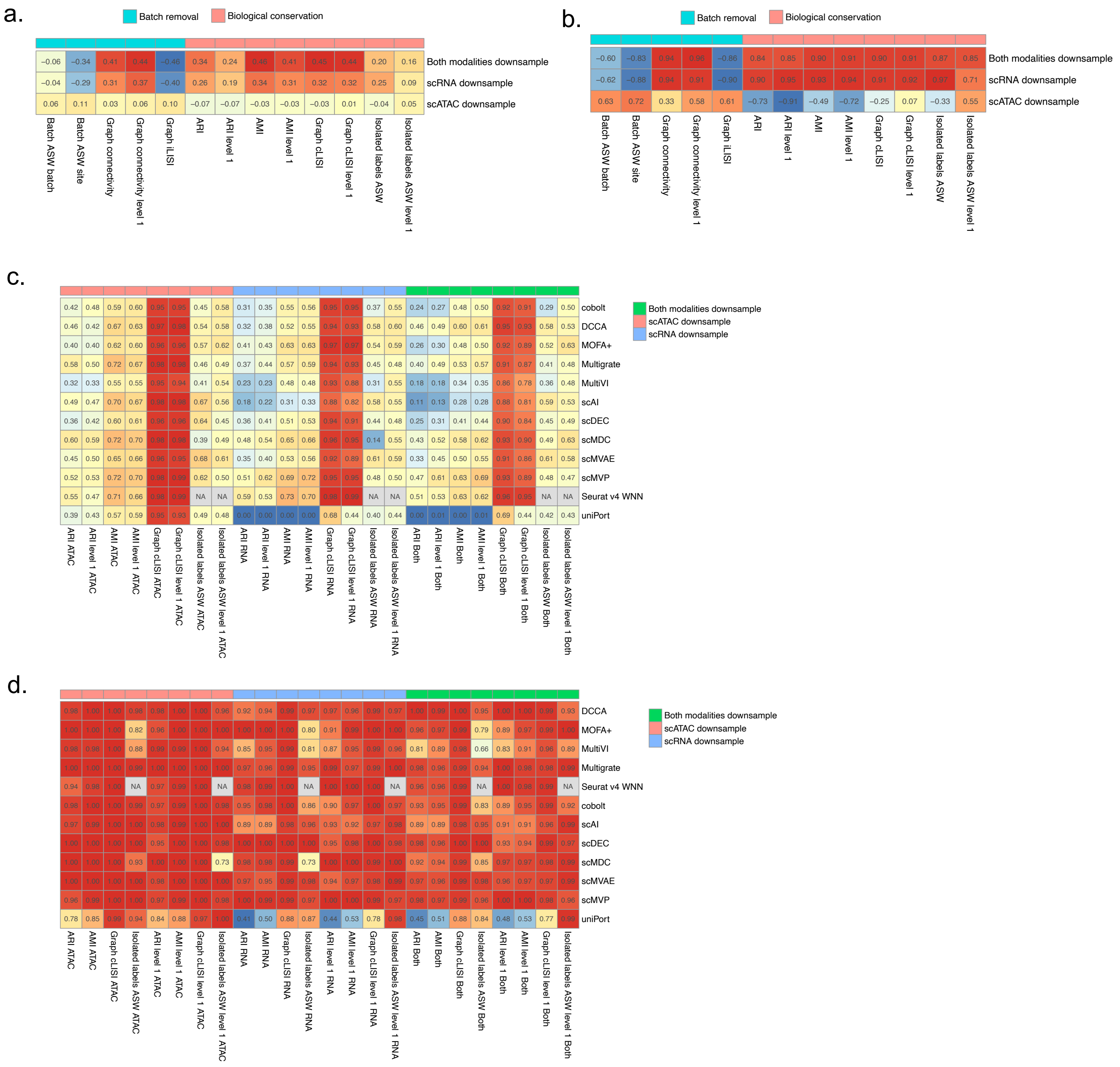

Supplementary Fig. 7. Details of the paired scRNA and scATAC integration robustness metrics in Fig. 2c and Extended Data Fig. 4d. (**a-b**). Correlations between metric values and downsample gradients. The color and number in heatmaps shows Pearson's r correlation between (**a**) metric values or (**b**) average metric values with the downsampling gradients. (**c**) Detail metric values for all paired scRNA and scATAC integration algorithms for BMMC p10 dataset at the 10% sequencing depth. (**d**) Detail downsample AUC values for all paired scRNA and scATAC integration algorithms.

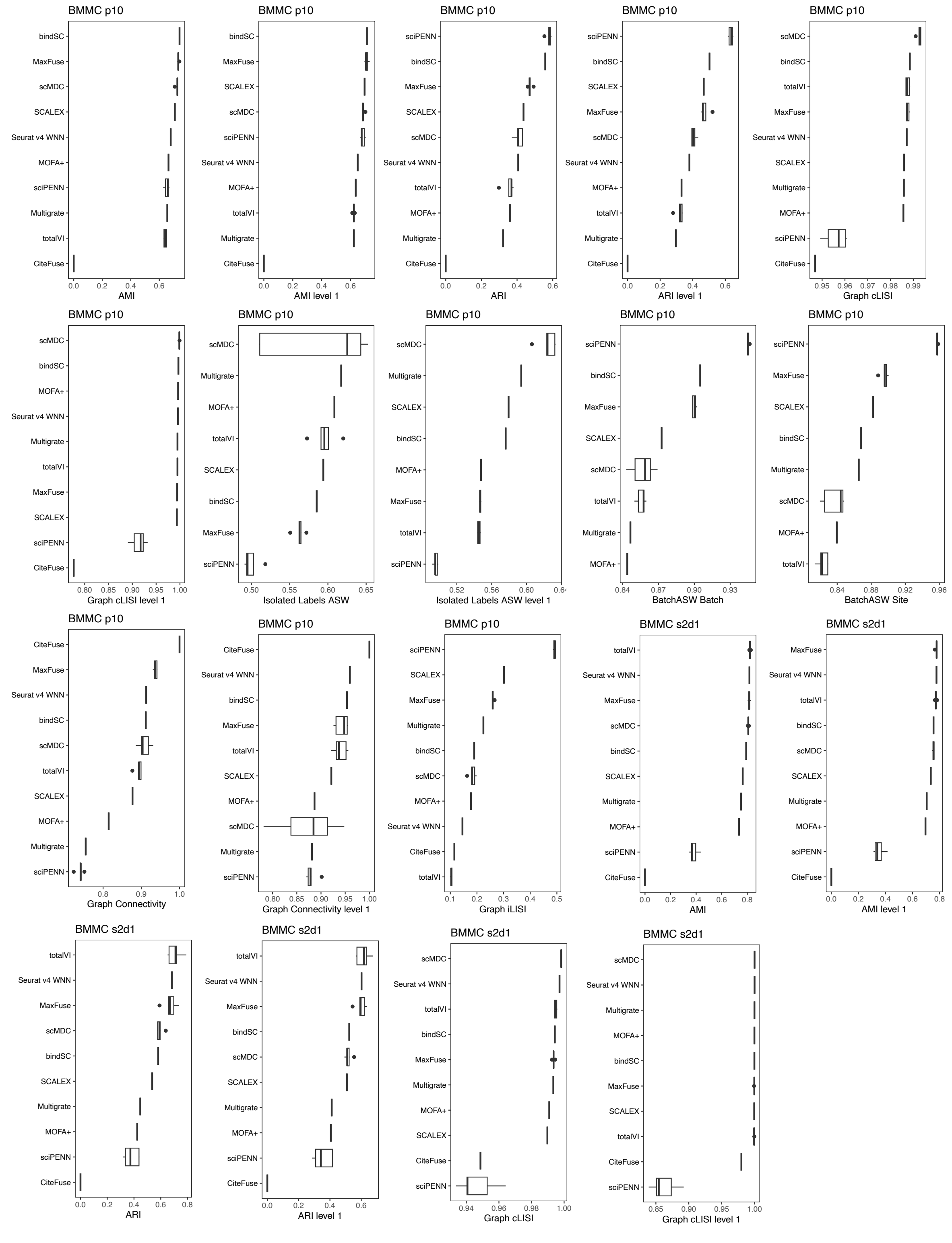


Supplementary Fig. 8. Boxplot of accuracy metrics for paired scRNA and ADT integration algorithms in BMMC CITE-seq p10 and s2d1 datasets, corresponding to Fig. 3b and Extended Data Fig. 5b. The dataset names are labelled at the top of each boxplot.

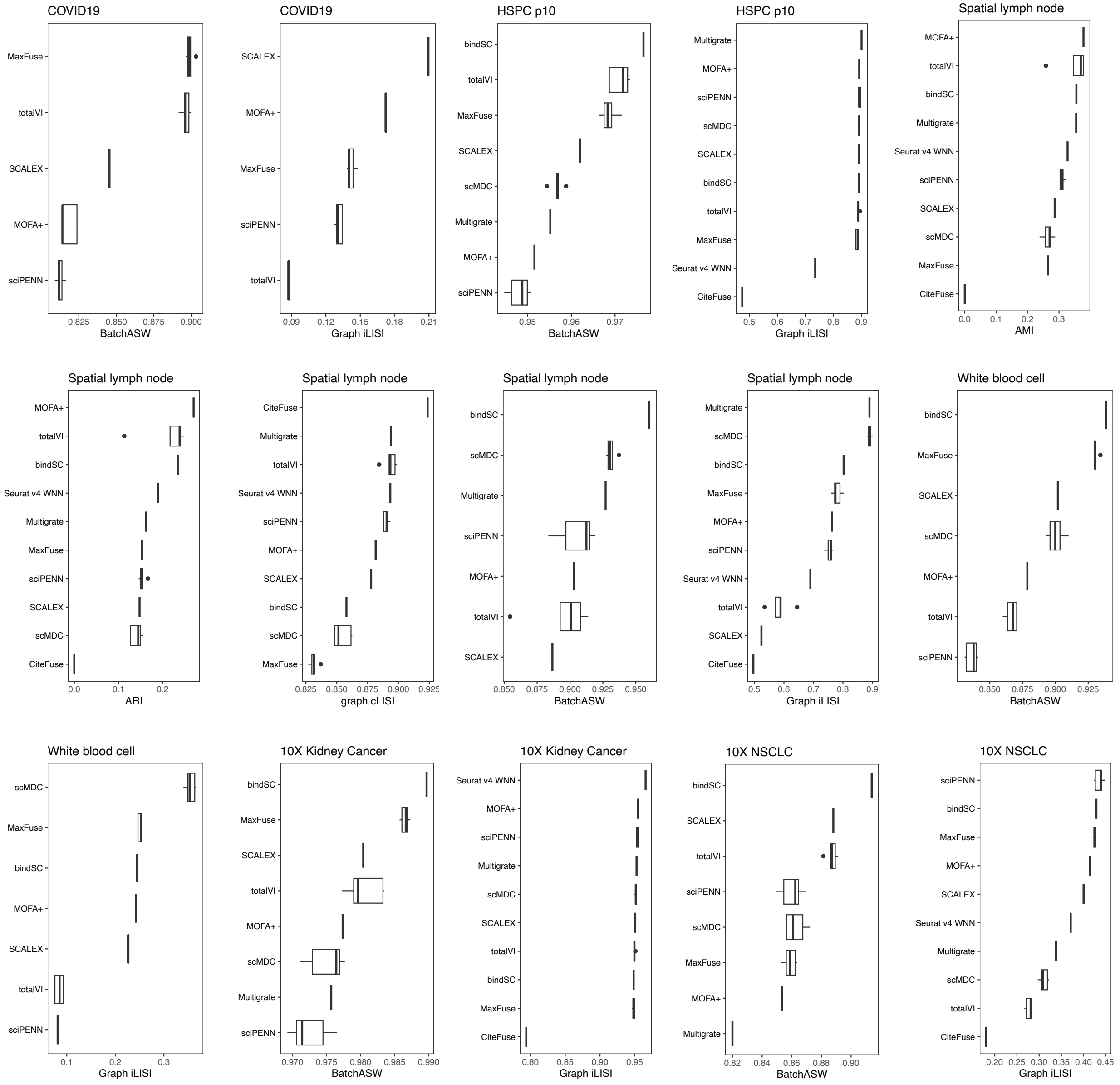


Supplementary Fig. 9 Boxplot of accuracy metrics for paired scRNA and ADT integration algorithms in other datasets except for BMMC CITE-seq datasets, corresponding to Fig. 3b and Extended Data Fig. 5b. The dataset names are labelled at the top of each boxplot.

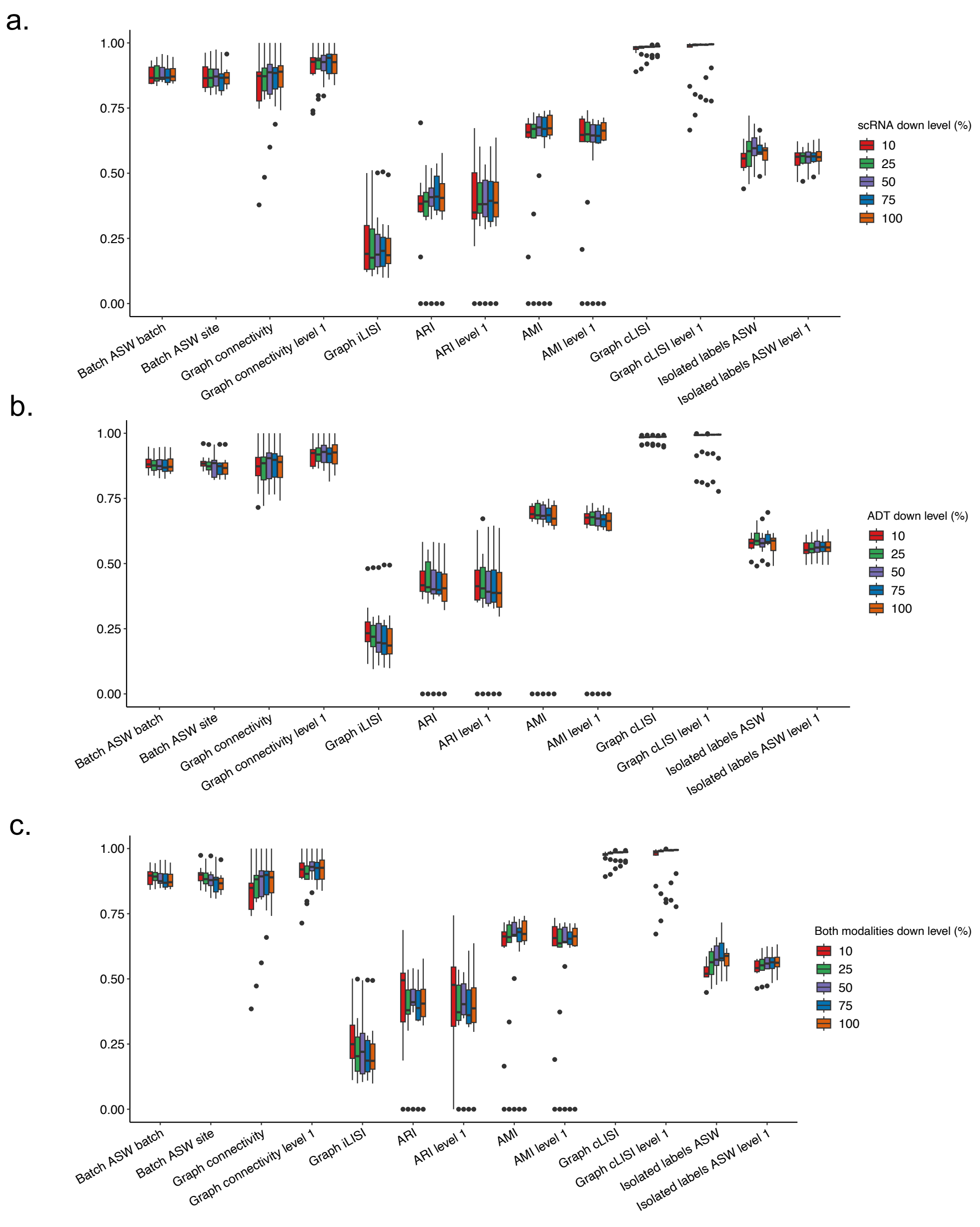


Supplementary Fig. 10. Boxplot of the biological conservation and batch removal metric values for all paired scRNA and ADT integration methods at all downsampling levels. This figure shows the metric scores distribution for simulations of **(a)** scRNA modality downsampling, **(b)** ADT modality downsampling and **(c)** both modalities downsampling.

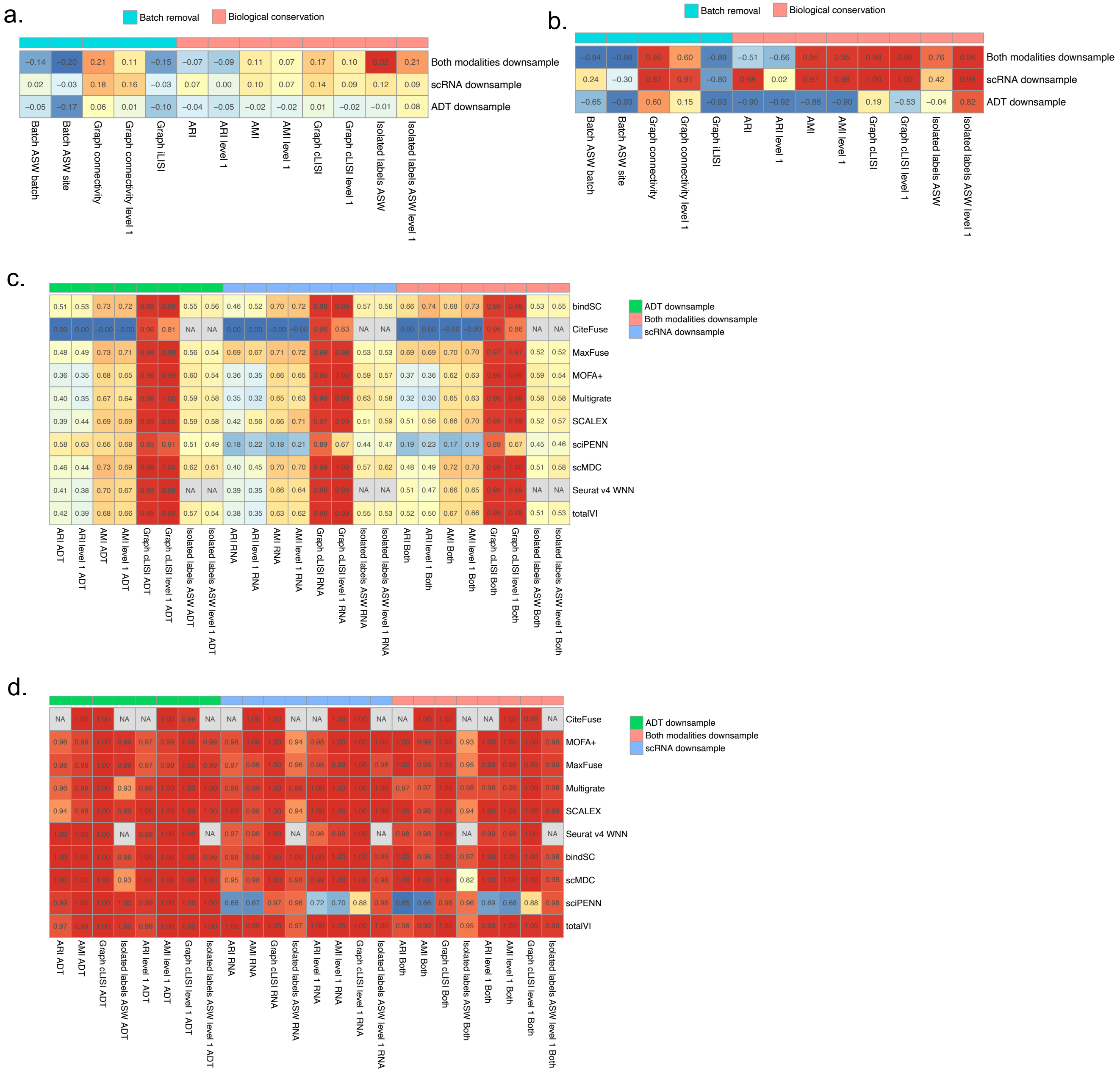


Supplementary Fig. 11. Details of the paired scRNA and ADT integration robustness metrics in Fig. 3d and Extended Data Fig. 5c. (**a-b**). Correlations between metric values and downsample gradients. The color and number in heatmaps shows Pearson's r correlation between (**a**) metric values or (**b**) average metric values with the downsampling gradients. (**c**) Detail metric values for all paired scRNA and ADT integration algorithms for BMMC p10 dataset at the 10% sequencing depth. (**d**) Detail downsample AUC values for all paired scRNA and ADT integration algorithms.

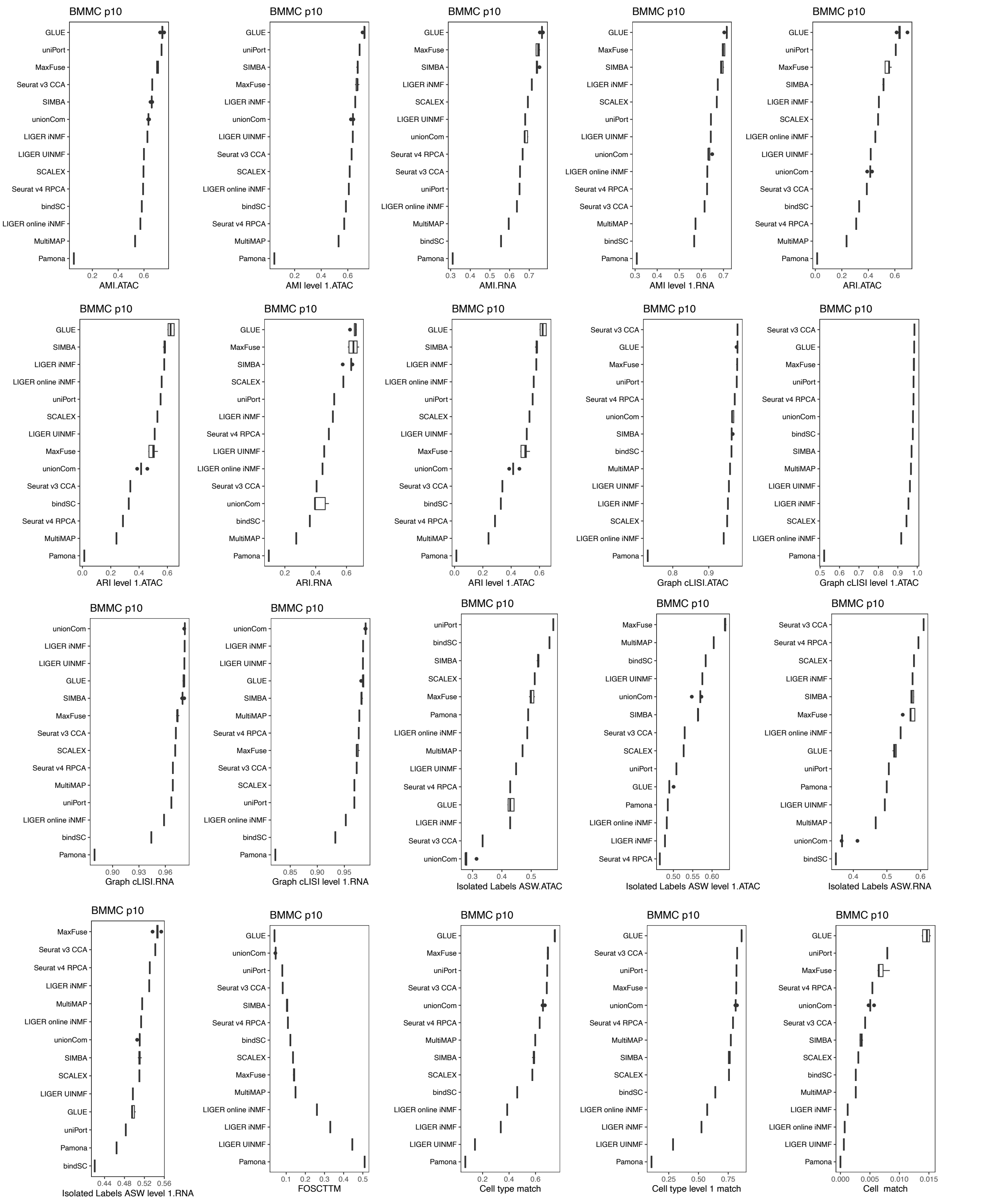


Supplementary Fig.12. Boxplot of accuracy metrics for unpaired scRNA and scATAC diagonal integration algorithms in BMMC Multiome p10 dataset, corresponding to Fig. 4c and Extended Data Fig. 6b.

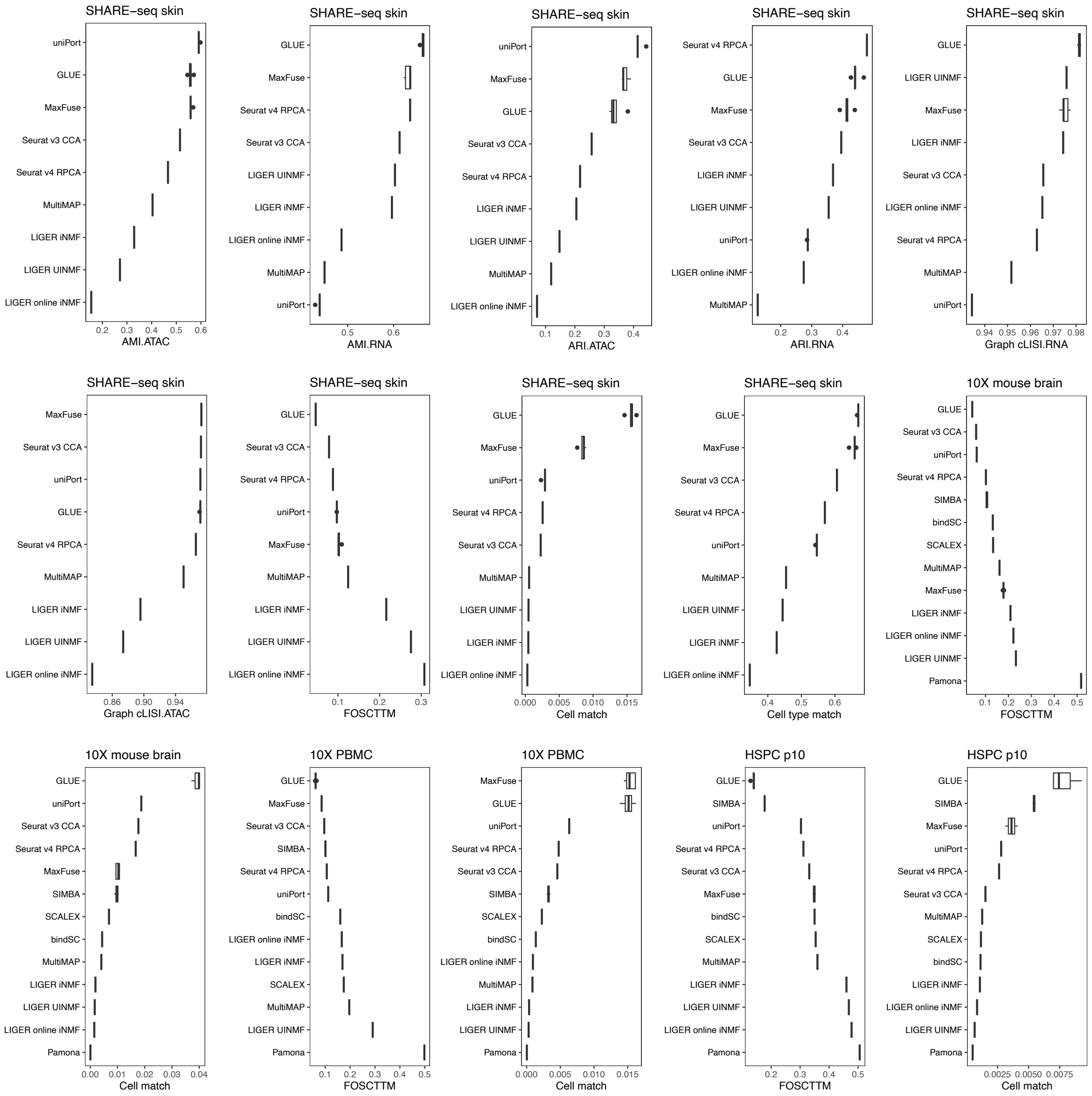


Supplementary Fig. 13. Boxplot of accuracy metrics for unpaired scRNA and scATAC diagonal integration algorithms in other datasets except for BMMC Multiome p10 dataset, corresponding to Fig. 4c and Extended Data Fig. 6b. The dataset names are labelled at the top of each boxplot.

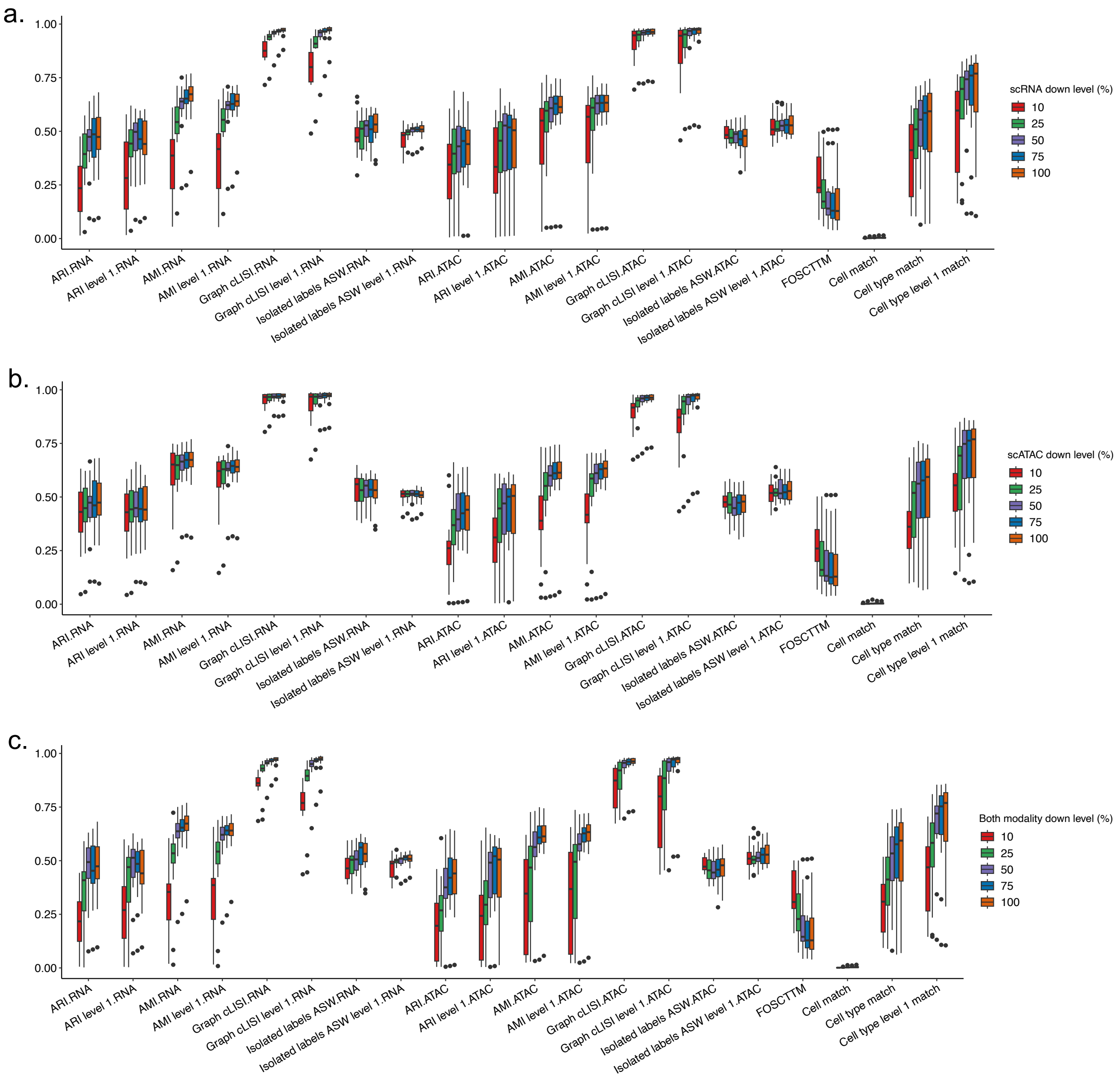


Supplementary Fig. 14. Boxplot of the embedding accuracy and cell alignment accuracy metric values for all unpaired scRNA and scATAC diagonal integration methods at all downsampling levels. This figure shows the metric scores distribution for simulations of **(a)** scRNA modality downsampling, **(b)** scATAC modality downsampling and **(c)** both modalities downsampling.

a.

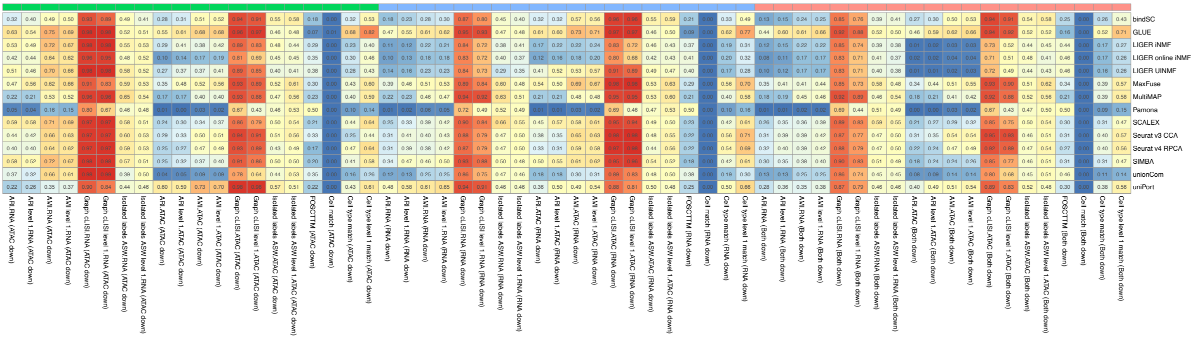


b.

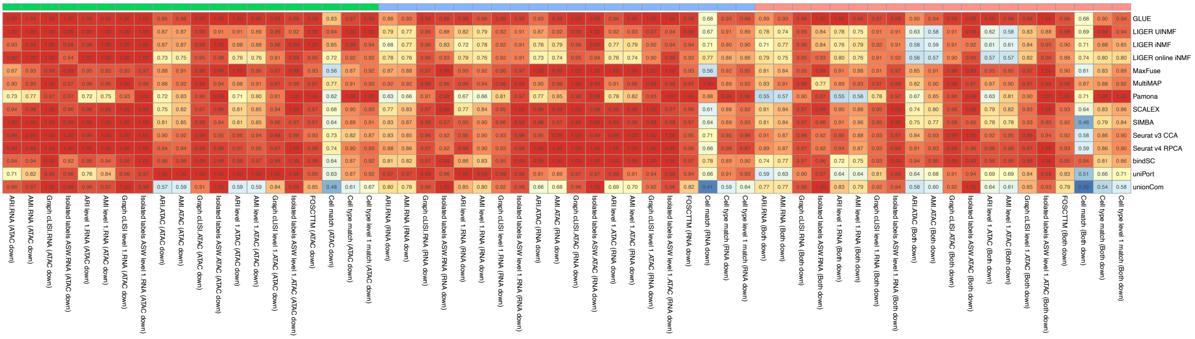


Supplementary Fig. 15. Details of the unpaired scRNA and scATAC diagonal robustness metrics in Fig. 4f and Extended Data Fig. 6c. (**a**) Detail metric values for all unpaired scRNA and scATAC diagonal integration algorithms for BMMC p10 dataset at the 10% sequencing depth. (**b**) Detail downsample AUC values for all unpaired scRNA and scATAC diagonal integration algorithms.

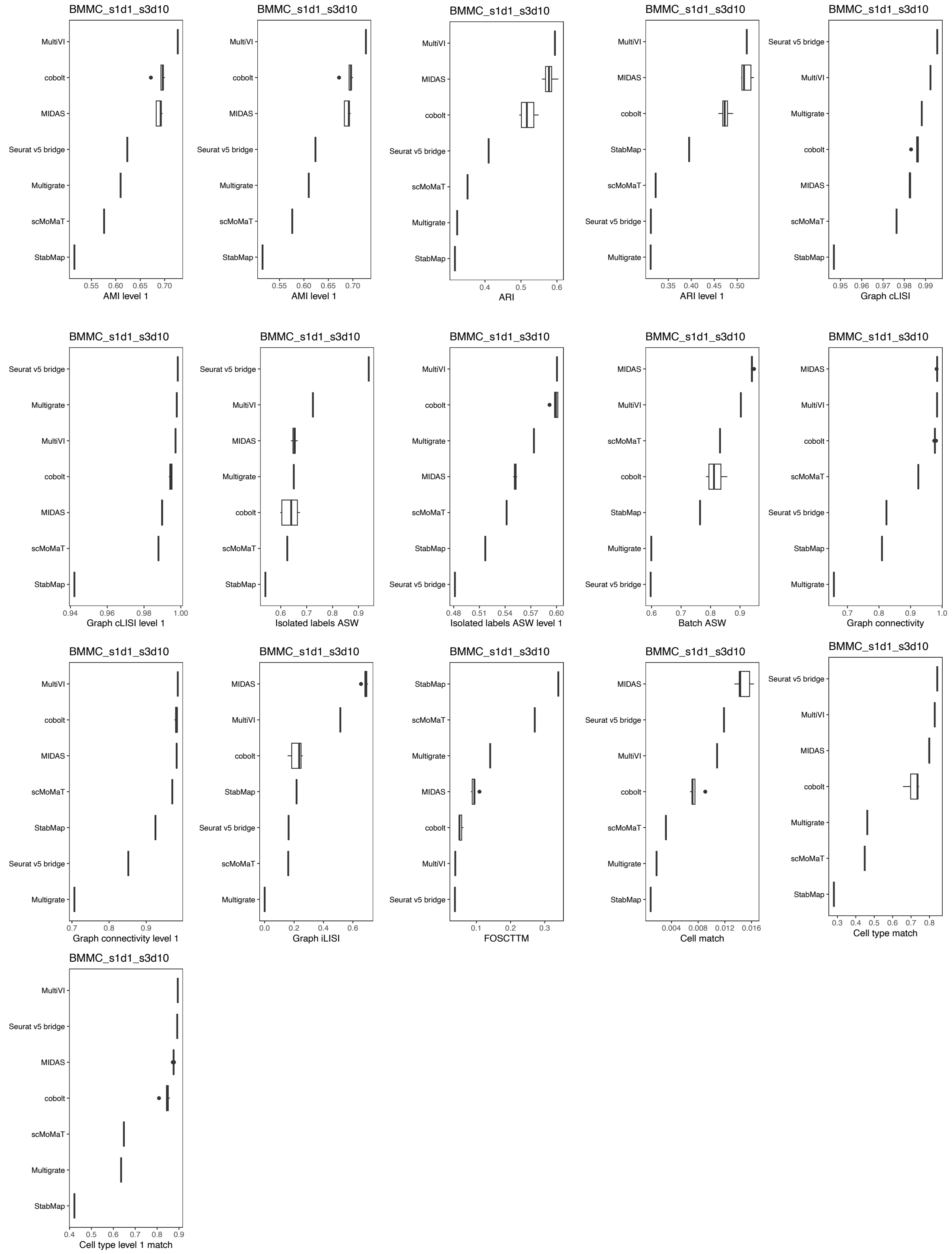


Supplementary Fig.16. Boxplot of the embedding accuracy and cell alignment accuracy metrics for unpaired scRNA and scATAC mosaic integration algorithms in BMMC Multiome s1d1_s3d10 dataset, corresponding to Fig. 5b and Extended Data Fig. 7b.

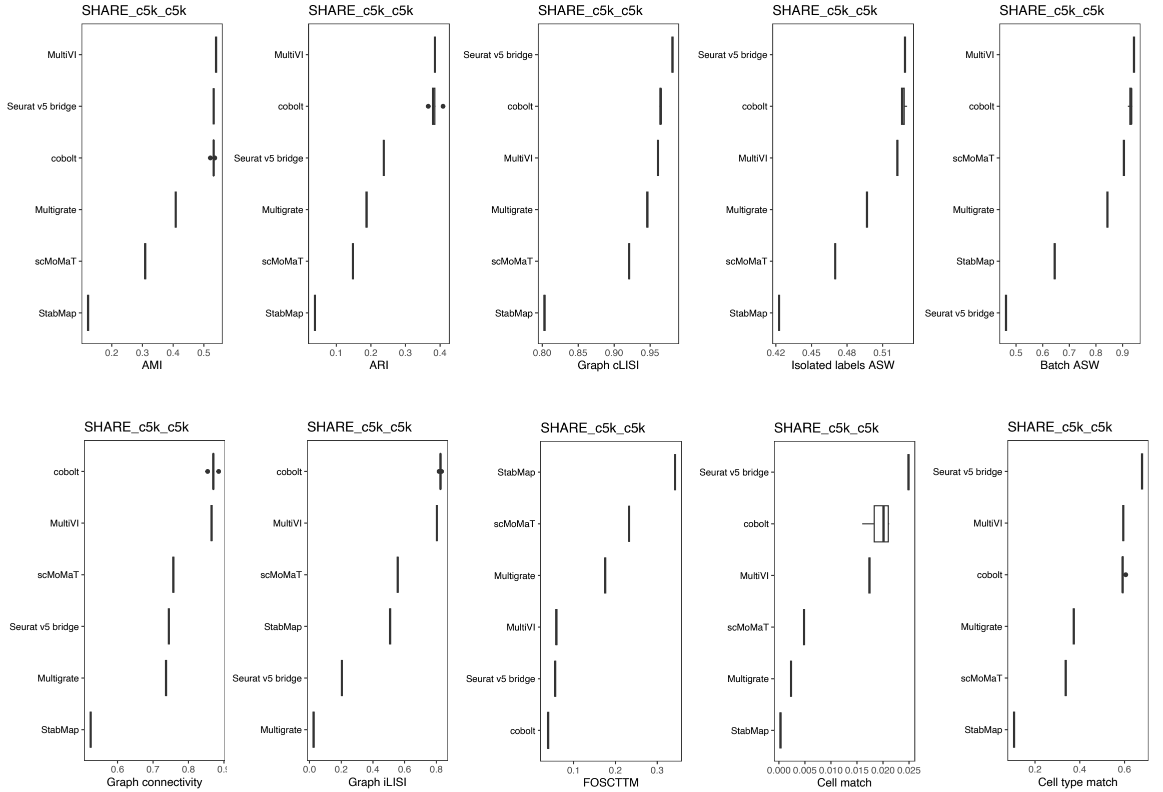


Supplementary Fig.17. Boxplot of the embedding accuracy and cell alignment accuracy metrics for unpaired scRNA and scATAC mosaic integration algorithms in SHARE-seq skin c5k_c5k dataset, corresponding to Fig. 5b and Extended Data Fig. 7b.

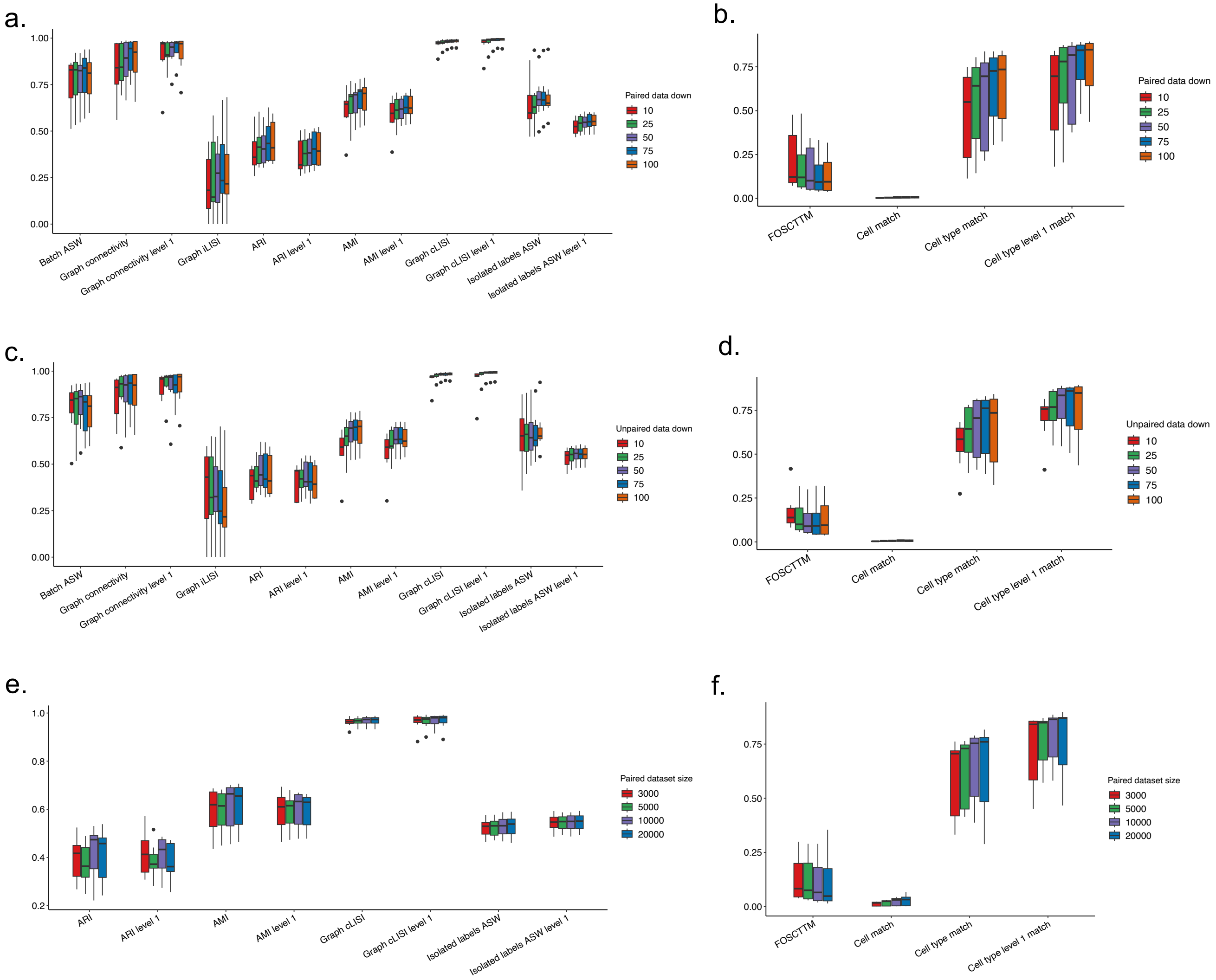


Supplementary Fig. 18. Boxplot of the embedding accuracy and cell alignment accuracy metric values for all unpaired scRNA and scATAC mosaic integration methods at (**a-b**) paired dataset downsampling levels, (**c-d**) unpaired dataset downsampling levels and (**e-f**) paired dataset sizes. The embedding accuracy and cell alignment accuracy metrics of different paired dataset sizes in (**e**) and (**f**) were calculated with unpaired cells only.

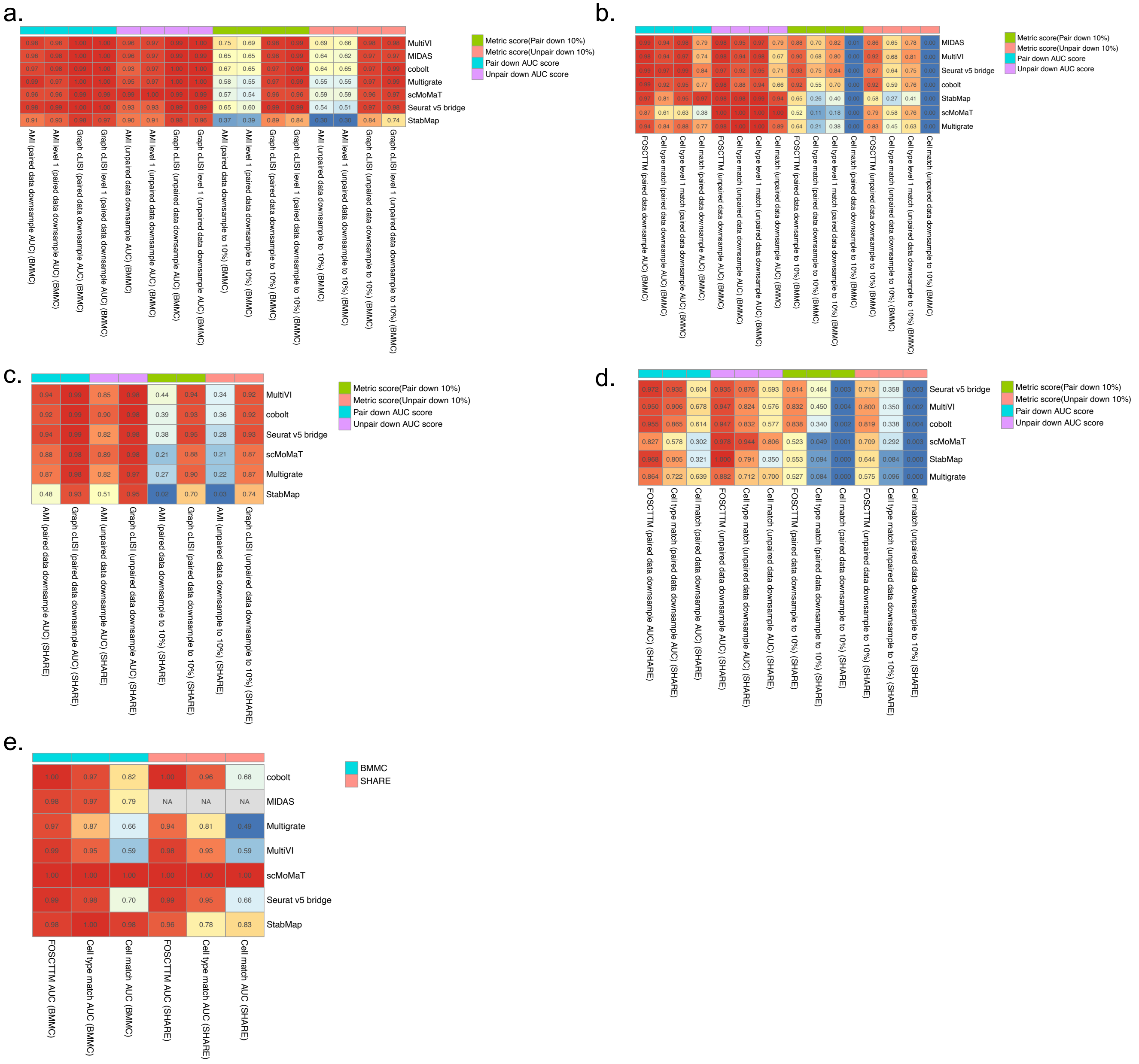


Supplementary Fig. 19. Details of the unpaired scRNA and scATAC mosaic integration robustness metrics in Fig. 5e. (**a-b**). Detail robustness scores of (**a**) embedding accuracy metrics and (**b**) cell alignment accuracy metrics for unpaired scRNA and scATAC mosaic integration algorithms in downsampling paired and unpaired multi-omic cells of BMMC Multiome s1d1_s3d10 dataset. (**c-d**). Detail robustness scores of embedding accuracy metrics and cell alignment accuracy metrics for all unpaired scRNA and scATAC mosaic integration algorithms in downsampling paired and unpaired multi-omic cells for SHARE-seq skin c5k_c5k dataset. (**e**) Detail AUCs of unpaired dataset cell alignment metrics in gradient paired dataset sizes for both BMMC and SHARE-seq datasets.

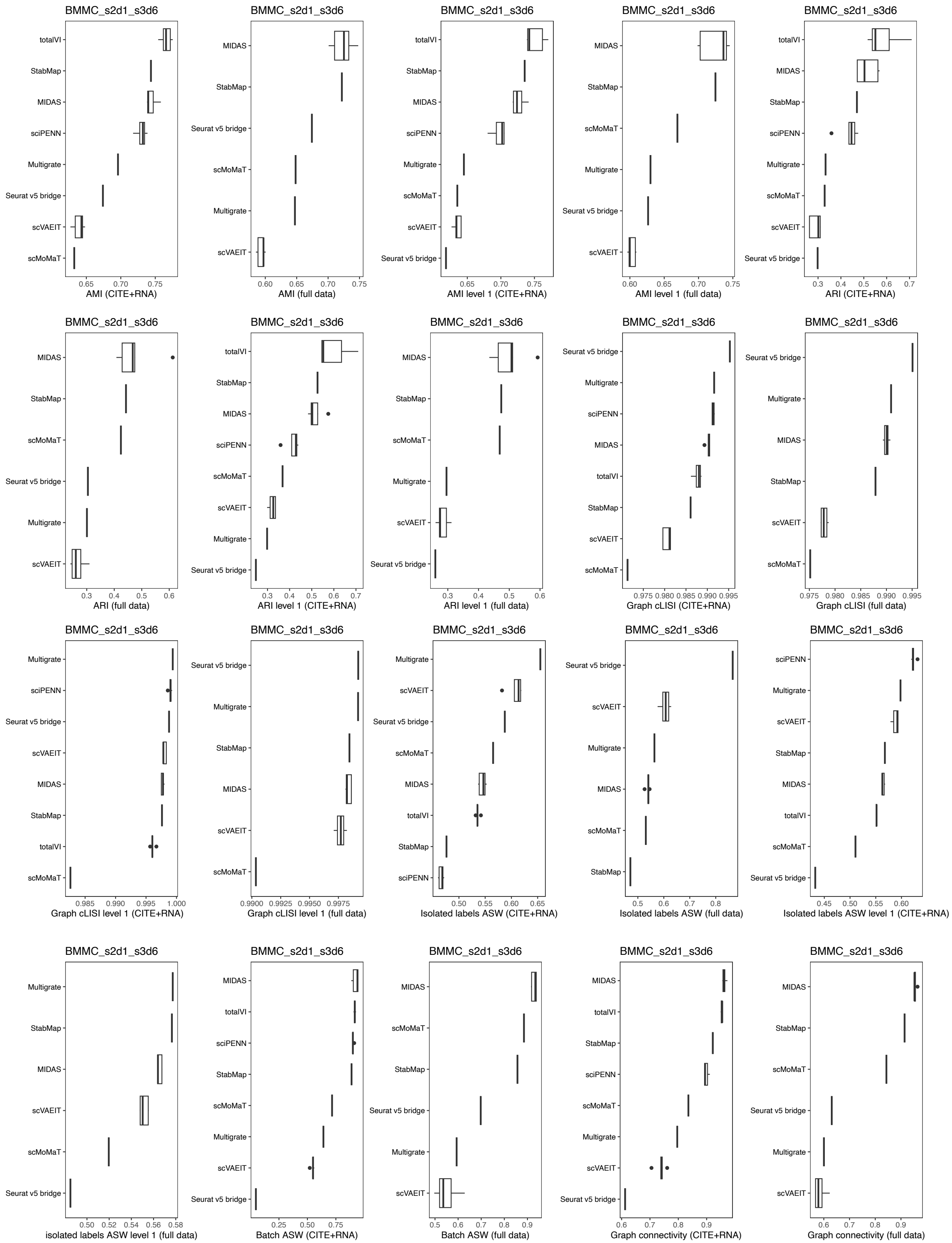

BMMC_s2d1_s3d6
totalVI
StabMap
MIDAS
sciPENN
Multigrate
Seurat v5 bridge
scVAEIT
scMoMaT
0.65 0.70 0.75
AMI (CITE+RNA)
BMMC_s2d1_s3d6
MIDAS
StabMap
Seurat v5 bridge
scMoMaT
Multigrate
scVAEIT
0.60 0.65 0.70 0.75
AMI (full data)
BMMC_s2d1_s3d6
totalVI
StabMap
MIDAS
sciPENN
Multigrate
scMoMaT
scVAEIT
Seurat v5 bridge
0.65 0.70 0.75
AMI level 1 (CITE+RNA)
BMMC_s2d1_s3d6
MIDAS
StabMap
scMoMaT
Multigrate
Seurat v5 bridge
scVAEIT
0.60 0.65 0.70 0.75
AMI level 1 (full data)
BMMC_s2d1_s3d6
totalVI
MIDAS
StabMap
sciPENN
Multigrate
scMoMaT
scVAEIT
Seurat v5 bridge
0.3 0.4 0.5 0.6 0.7
ARI (CITE+RNA)
BMMC_s2d1_s3d6
MIDAS
StabMap
scMoMaT
Seurat v5 bridge
Multigrate
scVAEIT
0.3 0.4 0.5 0.6
ARI (full data)
BMMC_s2d1_s3d6
totalVI
StabMap
MIDAS
sciPENN
scMoMaT
scVAEIT
Multigrate
Seurat v5 bridge
0.3 0.4 0.5 0.6 0.7
ARI level 1 (CITE+RNA)
BMMC_s2d1_s3d6
MIDAS
StabMap
scMoMaT
Multigrate
scVAEIT
Seurat v5 bridge
0.3 0.4 0.5 0.6
ARI level 1 (full data)
BMMC_s2d1_s3d6
Seurat v5 bridge
Multigrate
sciPENN
MIDAS
totalVI
StabMap
scVAEIT
scMoMaT
0.975 0.980 0.985 0.990 0.995
Graph cLISI (CITE+RNA)
BMMC_s2d1_s3d6
Seurat v5 bridge
Multigrate
MIDAS
StabMap
scVAEIT
scMoMaT
0.975 0.980 0.985 0.990 0.995
Graph cLISI (full data)
BMMC_s2d1_s3d6
Multigrate
sciPENN
Seurat v5 bridge
scVAEIT
MIDAS
StabMap
totalVI
scMoMaT
0.985 0.990 0.995 1.000
Graph cLISI level 1 (CITE+RNA)
BMMC_s2d1_s3d6
Seurat v5 bridge
Multigrate
StabMap
MIDAS
scVAEIT
scMoMaT
0.9900 0.9925 0.9950 0.9975
Graph cLISI level 1 (full data)
BMMC_s2d1_s3d6
Multigrate
scVAEIT
Seurat v5 bridge
scMoMaT
MIDAS
totalVI
StabMap
sciPENN
0.50 0.55 0.60 0.65
Isolated labels ASW (CITE+RNA)
BMMC_s2d1_s3d6
Seurat v5 bridge
scVAEIT
Multigrate
MIDAS
scMoMaT
StabMap
0.5 0.6 0.7 0.8
Isolated labels ASW (full data)
BMMC_s2d1_s3d6
sciPENN
Multigrate
scVAEIT
StabMap
MIDAS
totalVI
scMoMaT
Seurat v5 bridge
0.45 0.50 0.55 0.60
Isolated labels ASW level 1 (CITE+RNA)
BMMC_s2d1_s3d6
Multigrate
StabMap
MIDAS
scVAEIT
scMoMaT
Seurat v5 bridge
0.50 0.52 0.54 0.56 0.58
isolated labels ASW level 1 (full data)
BMMC_s2d1_s3d6
MIDAS
totalVI
sciPENN
StabMap
scMoMaT
Multigrate
scVAEIT
Seurat v5 bridge
0.25 0.50 0.75
Batch ASW (CITE+RNA)
BMMC_s2d1_s3d6
MIDAS
scMoMaT
StabMap
Seurat v5 bridge
Multigrate
scVAEIT
0.5 0.6 0.7 0.8 0.9
Batch ASW (full data)
BMMC_s2d1_s3d6
MIDAS
totalVI
StabMap
sciPENN
scMoMaT
Multigrate
scVAEIT
Seurat v5 bridge
0.6 0.7 0.8 0.9
Graph connectivity (CITE+RNA)
BMMC_s2d1_s3d6
MIDAS
StabMap
scMoMaT
Seurat v5 bridge
Multigrate
scVAEIT
0.6 0.7 0.8 0.9
Graph connectivity (full data)

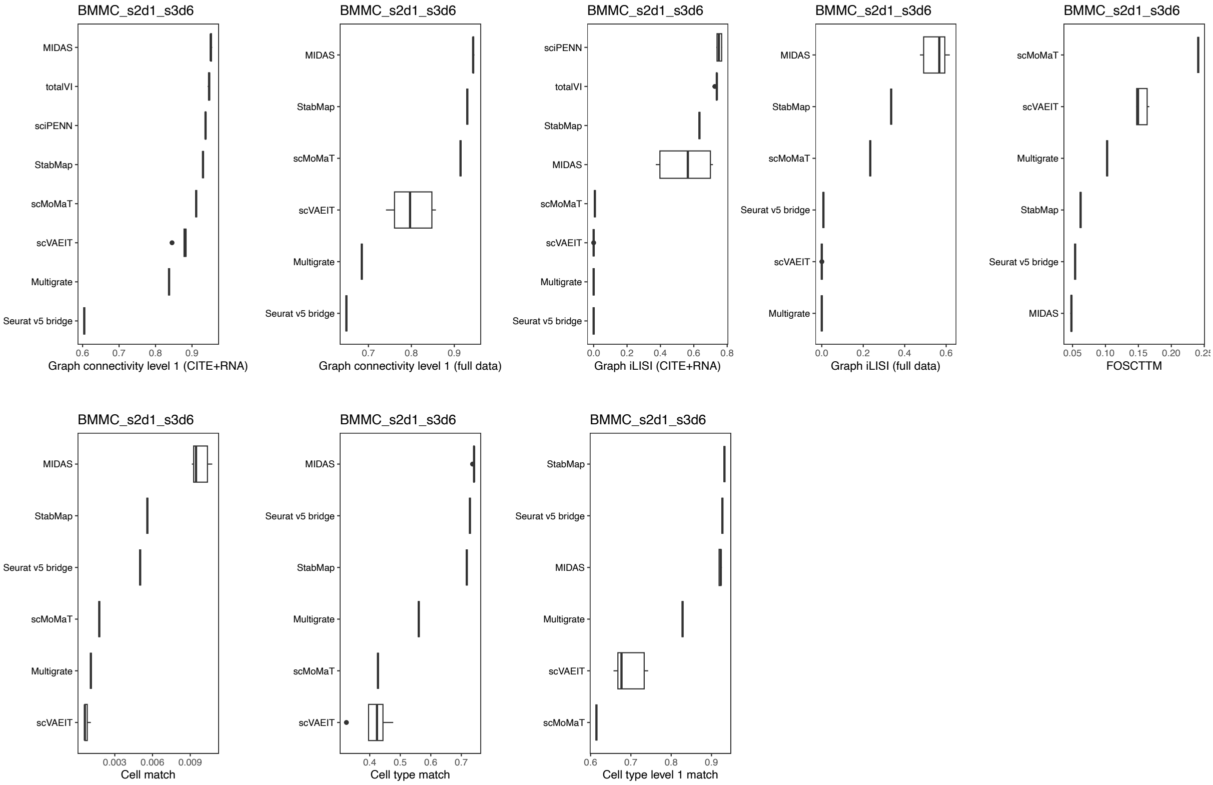


Supplementary Fig. 20. Boxplot of the embedding accuracy and cell alignment accuracy metrics for unpaired scRNA and ADT mosaic integration algorithms in BMMC CITE-seq s2d1_s3d6 dataset, corresponding to Fig. 6b and Extended Data Fig. 8b.

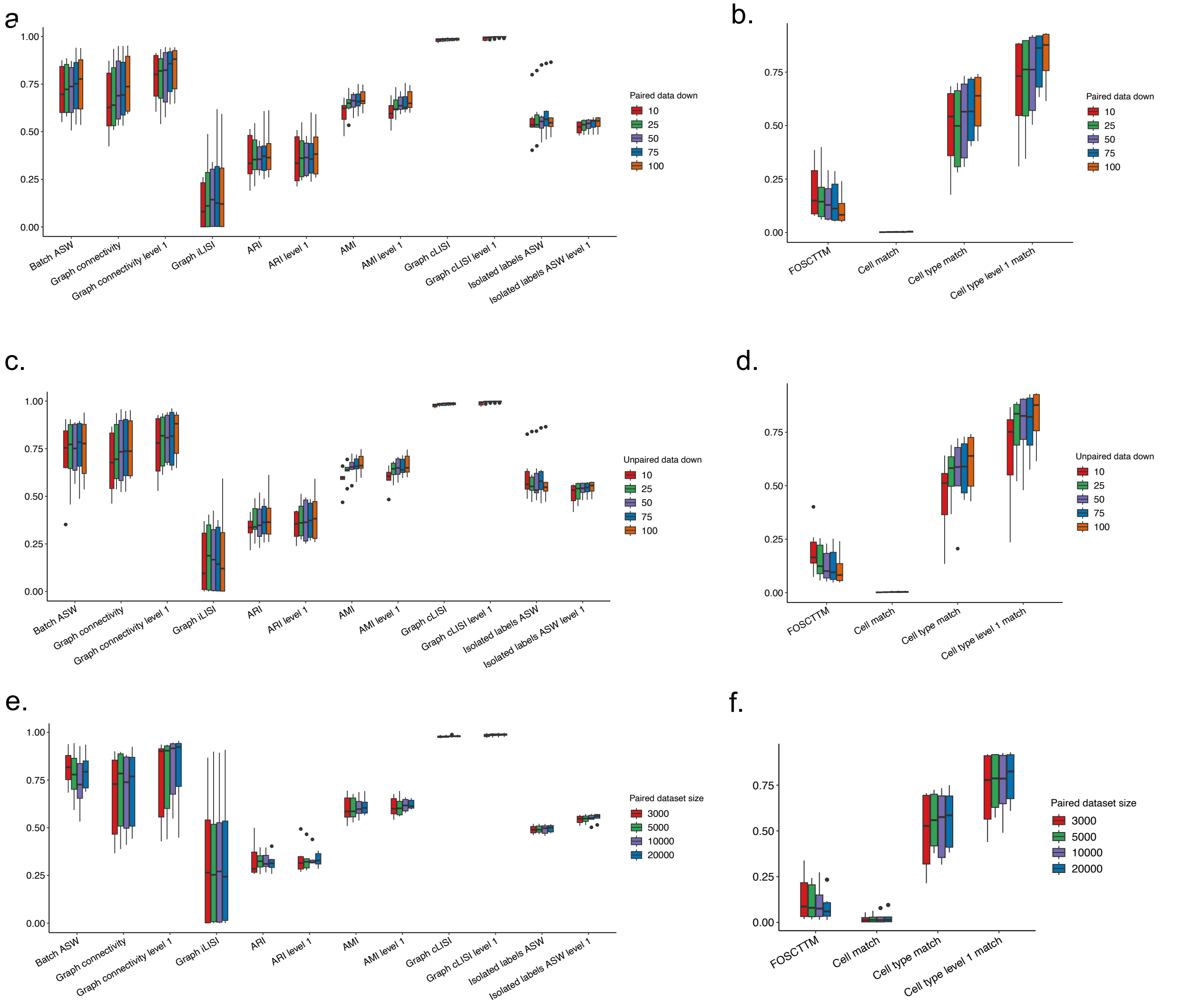


Supplementary Fig. 21. Boxplot of the embedding accuracy and cell alignment accuracy metric values for all unpaired scRNA and ADT mosaic integration methods at (**a-b**) paired dataset downsampling levels, (**c-d**) unpaired dataset downsampling levels and (**e-f**) paired dataset sizes. The embedding accuracy and cell alignment accuracy metrics of different paired dataset size in (**e**) and (**f**) were calculated with unpaired cells only,

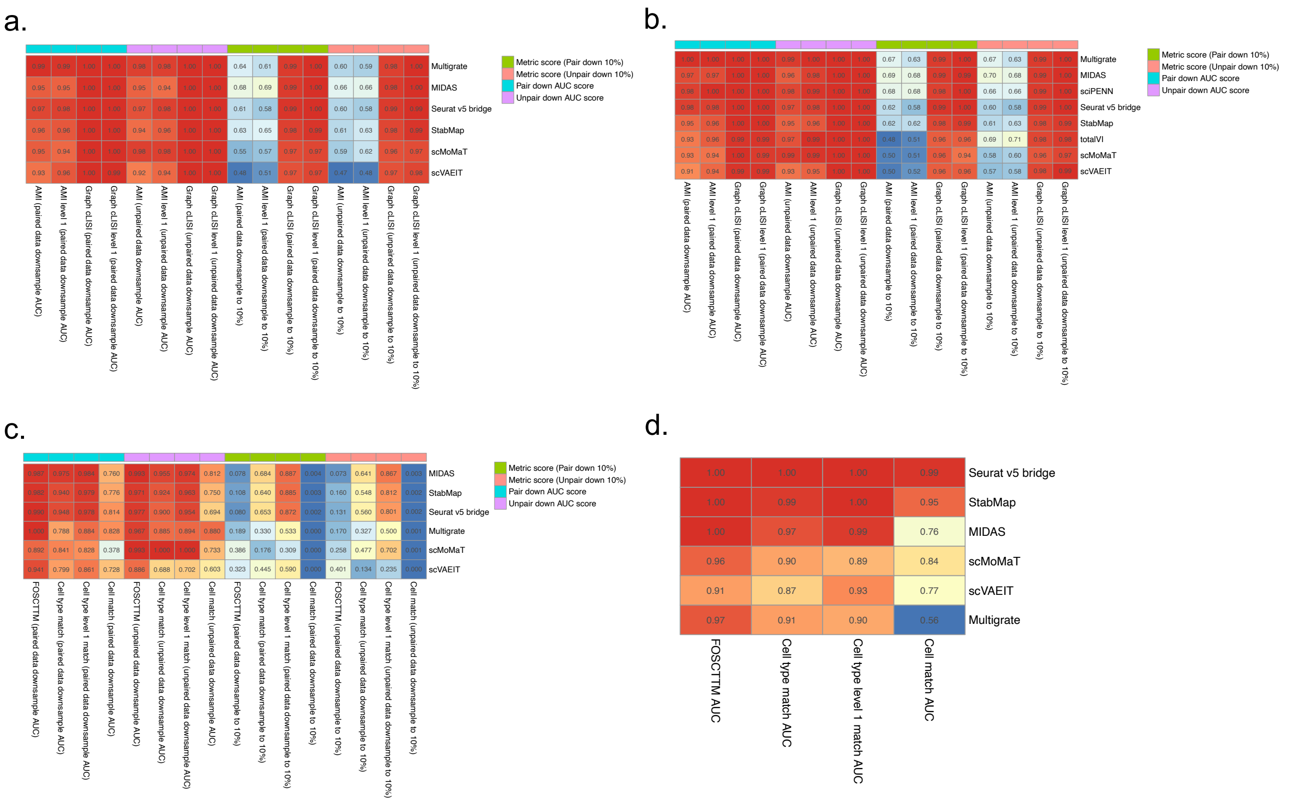


Supplementary Fig. 22. Details of the unpaired scRNA and ADT mosaic integration robustness metrics in Fig. 6c. (**a-b**). Detail robustness scores of embedding accuracy metrics of (**a**) full data and (**b**) CITE-seq and scRNA-seq cells in downsampling paired and unpaired multi-omic cells of BMMC CITE-seq dataset. (**c**) Detail robustness scores of cell alignment accuracy metrics cells in downsampling paired and unpaired multi-omic cells of BMMC s2d1_s3d6 dataset. (**d**) Detail AUCs of unpaired dataset cell alignment metrics in gradient paired dataset sizes of BMMC CITE-seq dataset.
